# A single-cell transcriptional roadmap of the mouse and human lymph node lymphatic vasculature

**DOI:** 10.1101/2019.12.31.892166

**Authors:** Menglan Xiang, Rubén Adrián Grosso, Akira Takeda, Junliang Pan, Tove Bekkhus, Kevin Brulois, Denis Dermadi, Sofia Nordling, Michael Vanlandewijck, Sirpa Jalkanen, Maria H. Ulvmar, Eugene C. Butcher

## Abstract

Single-cell transcriptomics promises to revolutionize our understanding of the vasculature. Emerging computational methods applied to high dimensional single cell data allow integration of results between samples and species, and illuminate the diversity and underlying developmental and architectural organization of cell populations. Here, we illustrate these methods in analysis of mouse lymph node (LN) lymphatic endothelial cells (LEC) at single cell resolution. Clustering identifies five well-delineated subsets, including two medullary sinus subsets not recognized previously as distinct. Nearest neighbor alignments in trajectory space position the major subsets in a sequence that recapitulates known and suggests novel features of LN lymphatic organization, providing a transcriptional map of the lymphatic endothelial niches and of the transitions between them. Differences in gene expression reveal specialized programs for (1) subcapsular ceiling endothelial interactions with the capsule connective tissue and cells, (2) subcapsular floor regulation of lymph borne cell entry into the LN parenchyma and antigen presentation, and (3) medullary subset specialization for pathogen interactions and LN remodeling. LEC of the subcapsular sinus floor and medulla, which represent major sites of cell entry and exit from the LN parenchyma respectively, respond robustly to oxazolone inflammation challenge with enriched signaling pathways that converge on both innate and adaptive immune responses. Integration of mouse and human single-cell profiles reveals a conserved cross-species pattern of lymphatic vascular niches and gene expression, as well as specialized human subsets and genes unique to each species. The examples provided demonstrate the power of single-cell analysis in elucidating endothelial cell heterogeneity, vascular organization and endothelial cell responses. We discuss the findings from the perspective of LEC functions in relation to niche formations in the unique stromal and highly immunological environment of the LN.

**Highlights**

Computational alignments (“trajectories”) predict LN LEC organization *in situ*, revealing a continuum of phenotypes punctuated by specialized clusters

Multiple intermediate phenotypes suggest LEC malleability

Gene profiles define niche-specific functional specialization

Medullary sinus LECs are comprised of Ptx3-LECs and Marco-LECs

- Distinct mechanisms for pathogen interactions and matrix modeling
- Ptx3-LECs: paracortical and central medullary sinuses near hilus; enriched for genes driving lymphangiogenic responses and lymphocyte egress
- Marco-LECs: peri-follicular medullary sinuses; macrophage-associated genes, complement and coagulation cascade

Niche-specific responses to inflammation

- IFN gene responses in SCS floor and medullary sinus LECs
- Suppression of LEC identity genes in responding subsets

Conserved and unique LEC subsets and gene programs across species

- Core subsets common to mouse and human
- Greater diversity of subsets and intermediates in human LN LECs

## Introduction

Lymph nodes (LNs) serve as hubs for the interaction and communication between tissue-derived and blood-derived immune cells [1]. Integrated along the large collecting lymphatic vessels, they are vital sensors of tissue damage, constantly sampling the incoming lymph [2]. The LN comprises a complex network of lymphatic sinuses surrounding a dense parenchyma, which mainly consists of immune cells but also specialized blood vessels and a network of mesenchymal cells [1; 2; 3]. Segregated B-cell (cortex) and T-cell (paracortex) areas characterize the LN architecture [4]. It is well established that the LN stromal cells play a central role in maintaining both this structure and the immunological functions of the LN, providing chemotactic cues, cytokines and a structural reticular framework that guide immune cell positioning, migration, survival and activation (reviews: [3; 5]). Single-cell sequencing has enabled delineation of nine distinct clusters of murine LN mesenchymal cell phenotypes [6] underlining the complexity needed to maintain the LN structure and coordinate immunity.

The lymphatic vasculature is the first structural component of LN encountered by incoming lymph-borne molecules or cells. Recent studies have revealed an intriguing regional specialization and cellular heterogeneity that characterize the LN lymphatic endothelium and differentiate the LN lymphatic endothelial cells (LECs) from LECs in peripheral lymphatic vessels [7; 8; 9; 10; 11]. Subset specific markers with functional implications include the atypical chemokine receptor ACKR4 (also known as CCRL1), specifically expressed by the LEC layer that forms the ceiling (cLECs) of the subcapsular sinus (SCS) [9], where lymph enters from the afferent collecting vessels. ACKR4 is a scavenger receptor for the homeostatic chemokines CCL19, 21 and 25 [12] and controls entry of tissue-derived dendritic cells (DCs) into the LN through controlling the formation of CCL21 chemokine gradients across the SCS [9]. Leukocyte entry occurs primarily through the SCS floor LECs (fLECs), which in mouse express the Mucosal vascular Addressin Cell Adhesion Molecule 1 (MAdCAM-1) [7; 13] among other adhesion and attractant molecules that can control leukocyte transmigration [7; 14]. The SCS also functions as a physical barrier and gateway, enabling size-restricted access of antigens to the LN parenchyma [15]: the glycoprotein plasmalemma vesicle-associated protein (PLVAP) together with cLEC-expressed Caveolin 1 (CAV1) [10], form sieve-like diaphragms in the transendothelial channels that bridge the SCS to the conduit system, and that descend from the SCS floor [10]. This structural barrier is complemented by a dense network of macrophages closely associated with SCS, providing essential innate immune functions and a filtering system for pathogens [16]. The SCS and medullary sinus macrophage niches in the LN were recently shown to be dependent on LEC expressed CSF-1 (Colony stimulating factor −1) [17] and Receptor activator of nuclear factor κ B (RANK)/RANKL signaling between the LN LECs and SCS lining mesenchymal cells [18].

LN LECs can also directly influence adaptive immune responses, either through presentation of tissue antigens, which contributes to the maintenance of peripheral tolerance [19; 20; 21], or by serving as reservoirs for antigens [22]. LN LECs express the immune-check-point ligand Programmed Death-Ligand 1 (PD-L1) (also known as CD274) [7; 23], an inhibitor of T cell activation, and lack expression of co-stimulatory genes [23], which may explain their role in tolerance. PD-L1 is expressed selectively in the floor of the SCS (fLECs), cortical sinuses and parts of the medulla [7; 23]. Genes that influence the communication between LECs and their surroundings could contribute to endothelial regulation as well, and interestingly, PD-L1 expression was recently shown to moderate proliferation and enhance survival of LN LECs in inflammation [24]. The diverse and site-specific specialization of the LN lymphatic endothelium is at least partly dependent on cross-talk with immune cells, with contributions from B-cells, T-cells [7; 25] and mesenchymal stromal cells [18]. Hence, the LNs provide both a unique model system to explore endothelial cell interactions with their surroundings and a model for exploring endothelial diversity and phenotypic plasticity.

Our recent single-cell analysis of the human LN LECs revealed the complexity and specialization of the LN lymphatic endothelium in man [26]. A detailed profiling of the mouse lymphatic endothelium and comparison of human and mouse LN lymphatic endothelium is still missing. Here we provide single-cell transcriptomic analysis of the mouse LN lymphatic vasculature. We show that computational alignments (relationships revealed by nearest neighbor trajectory inference) recapitulate key aspects of the tissue architecture and predict physical relationships between LN LECs in the tissue, illustrating the power of single-cell analysis for understanding the organization of the vascular endothelium. Cross-species analyses further allowed us to define conserved and divergent LEC phenotypes and lymphatic vascular niches. Notably, the analysis delineates two specialized subsets of medullary sinus LECs that are distinct in gene expression and location in the LN, present in both mouse and human. Using the mouse LN response to cutaneous oxazolone as a model, we show that sites of immune cell entry and exit from the LN, fLECs and medullary sinuses respectively, respond rapidly to inflammation, whereas structurally important cLECs are less affected. Together, the results demonstrate the power of bioinformatic tools for elucidating endothelial cell heterogeneity, physical relationships and cellular responses in complex vascular beds, and provide a basis for future detailed analysis of human and mouse LN LEC responses in disease.

## Results and discussion

### Single-cell trajectories model tissue architecture and physical relationships between LN LECs

To assess the heterogeneity of the lymphatic vasculature and the relationships between lymphatic vascular niches, we analyzed lin-Pdpn+CD31+ cells from mouse peripheral LNs (i.e. axillary, brachial and inguinal) by single-cell RNA sequencing (scRNA-seq) using the 10x Genomics system (Figure 1A). An independent set of LN LECs was sorted as single cells and subjected to SMART-seq2 analysis [27] (Figure 1A). *Prospero homeobox protein 1* (*Prox-1*) was used as a pan-LEC marker in the analysis [28]. Blood endothelial cells (*Flt1+Ly6c1*+), immune cells (CD45+) and pericytes (*Pdgfra+* or *Pdgfrb+*) were identified by marker gene expression and excluded from further analysis. We used a combination of unsupervised clustering and graph-based methods to determine LEC subsets (Figure 1B; and Methods).

**Figure 1:**
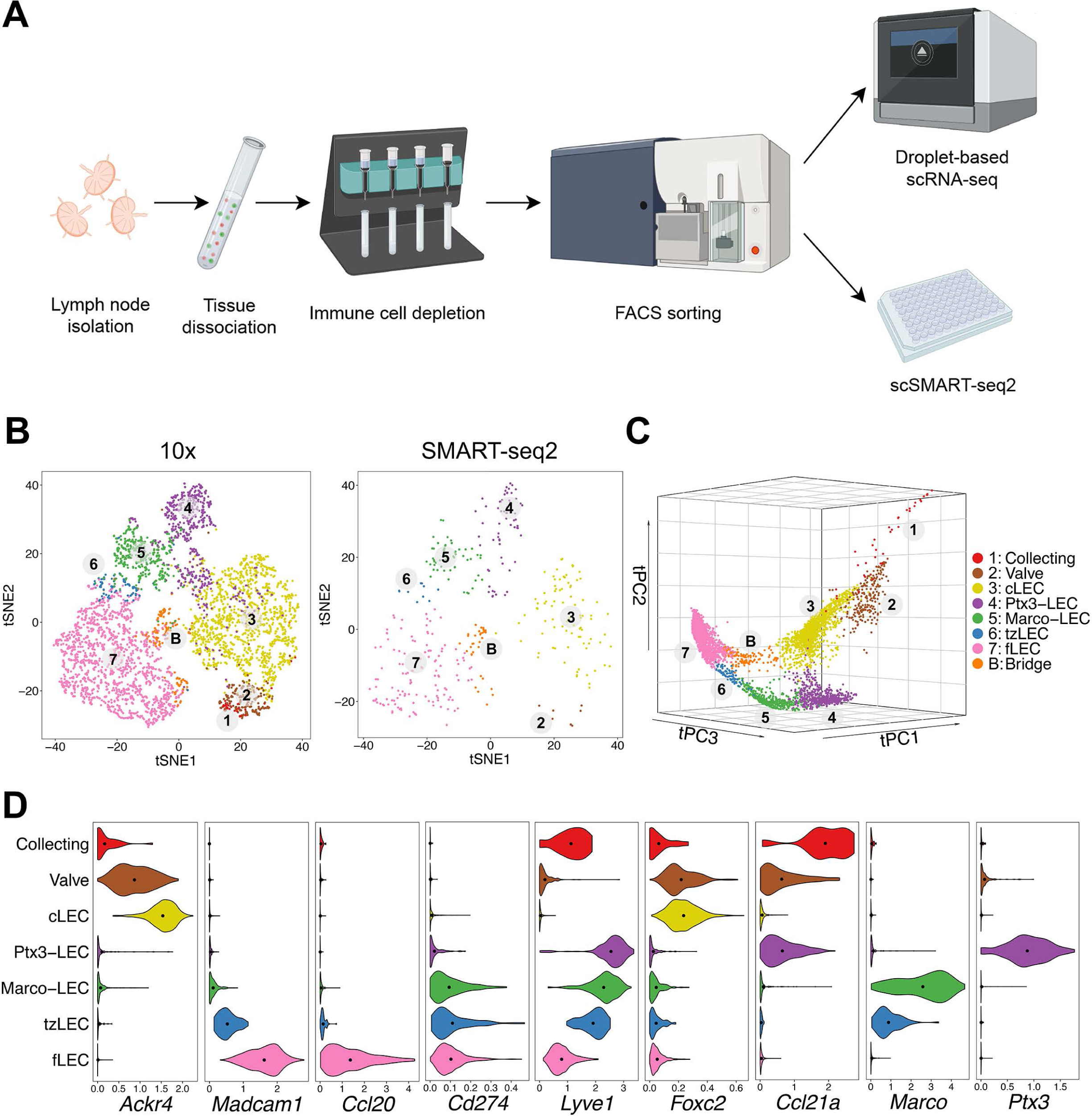
Mouse LN LEC subsets revealed by 10x and SMART-seq2. **(A)** Experimental workflow. Mouse peripheral LNs were isolated and dissociated into single-cell suspensions. Hematopoietic cells were MACS-depleted, and EC (lin-CD31+) were FACS-sorted for single-cell sequencing using the droplet-based 10x Genomics system or the SMART-seq2 workflow (figure created with BioRender.com). **(B)** tSNE plots of 4252 LECs processed by 10x (left) and 383 LECs by SMART-seq2 (right). Cells are colored by subset. Numbers indicate the continuum of the lymphatic endothelium. **(C)** tSpace analysis of single-cell trajectories depicting nearest neighbor alignments of LECs. **(D)** Expression of LEC subset defining genes in C57BL/6 mouse (10x). Dots indicate mean log-normalized transcript count.

In addition to gene profile-based dimensionality reduction with t-distributed Stochastic Neighbor Embedding (tSNE) (Figure 1B) and Uniform Manifold Approximation and Projection (UMAP) (see below), we used a trajectory analysis algorithm, tSpace [29] to define high dimensional nearest neighbor alignments that emphasize the continuum of cell phenotypes and reveal transitions between related cells (Figure 1C). In developing cell systems trajectory inference methods can model developmental sequence (“pseudotime”) [29]. In the resting adult vascular endothelium, they facilitate computational modeling of vessel architecture [30]. Subsets and alignments were shared in mice with different genetic backgrounds (C57BL/6 and BALB/c) (Figure S1A). Cells from both male and female mice were represented in the dataset and comprised similar phenotypes (Figure S1B).

Our analyses identified major clusters representing LN cLECs and fLECs based on the expression of known SCS ceiling marker *Ackr4* [9] as well as SCS floor markers *Madcam-1* [7; 13] and the chemokine *Ccl20* [14; 26] (Figures 1D and S1C). Candidate valve-related LECs display high expression of known lymphatic valve markers, including the transcription factor *Forkhead box protein C2* (*Foxc2*) [31] (Figures 1D and S1C). A minor cluster, most apparent in tSpace projections (Figure 1C) and associated with valve, expressed higher levels of peripheral lymphatic vessels markers including *Lymphatic vessel endothelial hyaluronan receptor 1* (*Lyve-1*) and the chemokine *Ccl21*, together with lower expression of *Foxc2* compared to the candidate valve LECs (Figure 1D). This population likely represents collector or pre-collector LECs [2; 31]. These subsets were poorly represented in our data and we discuss them in the context of the architecture of LN LECs, but exclude them from detailed differential gene expression analyses below. Unsupervised clusters separate *Lyve-1* high candidate medullary sinus LEC into two subsets, referred to here as Ptx3-LECs (*Ptx3*+) and Marco-LECs (*Marco*+) (Figures 1D and S1C).

Alignment of LECs in tSpace visualization recapitulates known connections within the complex lymphatic endothelial network in LN and reveals previously unappreciated relationships as well as intermediate, putative transitional phenotypes. In addition to the subsets identified by clustering, we highlight two transitional populations here, for discussion below. Transition zone tzLECs comprise a link between fLECs and Marco-LECs in tSpace (Figure 1C). Additionally, a minor “bridge” population (B) aligns along a direct path between fLECs and cLECs in tSpace projection (Figure 1C). As emphasized by numbering of subsets in Figure 1C and illustrated schematically in Figure 4A, trajectories starting from candidate collecting LECs (1) lead prominently to valve (2) and to SCS ceiling LEC (3), consistent with their known physical connections. cLECs branch to fLECs through the bridging population (B) but also transition prominently to Ptx3-LECs (4), Ptx3-LECs to Marco-LECs (5), and Marco-LECs via tzLECs (6) to fLEC (7) along a phenotypic sequence or path that is well represented in all LEC datasets here.

### *In situ* localization of LN LEC subsets

To define LEC subsets and their niches *in situ*, we carried out immunofluorescence staining using antibodies to subset differentiating markers predicted from gene expression (Figure 1D) in inguinal LNs of *Prox1-GFP* reporter mice [32], where all LECs express the fluorescent reporter GFP (Figure 2). scRNA-seq predicts that *Lyve-1* is absent in cLECs and low in fLECs, but is highly expressed in Marco-LECs and Ptx3-LECs (Figure 1D). Staining for LYVE-1 highlights the cortical, paracortical and entire medullary region of mouse inguinal LN (Figures 2A and 2B). Expression of *Cd274* (Figure 1D), which encodes the immune checkpoint inhibitor PD-L1, distinguishes fLECs, Marco-LECs and tzLECs (*Cd274*_*h*i_) from Ptx3-LECs (*Cd274*_*l*ow_) (Figures 1D and S1C). Co-immunostaining of PD-L1 and LYVE-1 shows that PD-L1 expression defines the fLECs and discrete regions of LYVE-1_high_ medullary sinus LECs that are distant from the hilus (Figure 2A). Expression of *Marco,* which encodes the scavenger receptor Macrophage Receptor with Collagenous Structure (MARCO), is highly selective for Marco-LECs (Figure 1D): MARCO is a known medullary sinus marker in human LN [26; 33]. We show that MARCO expression pattern in the mouse LN medulla selectively overlaps with that of PD-L1 (Figure 2B). Co-staining of MARCO and the B-cell marker B220 shows that MARCO_high_ regions start from the cortical areas (defined based on presence of B-cell follicles) and extend into the peri-follicular medulla of both inguinal and popliteal LNs (Figure S2).

**Figure 2:**
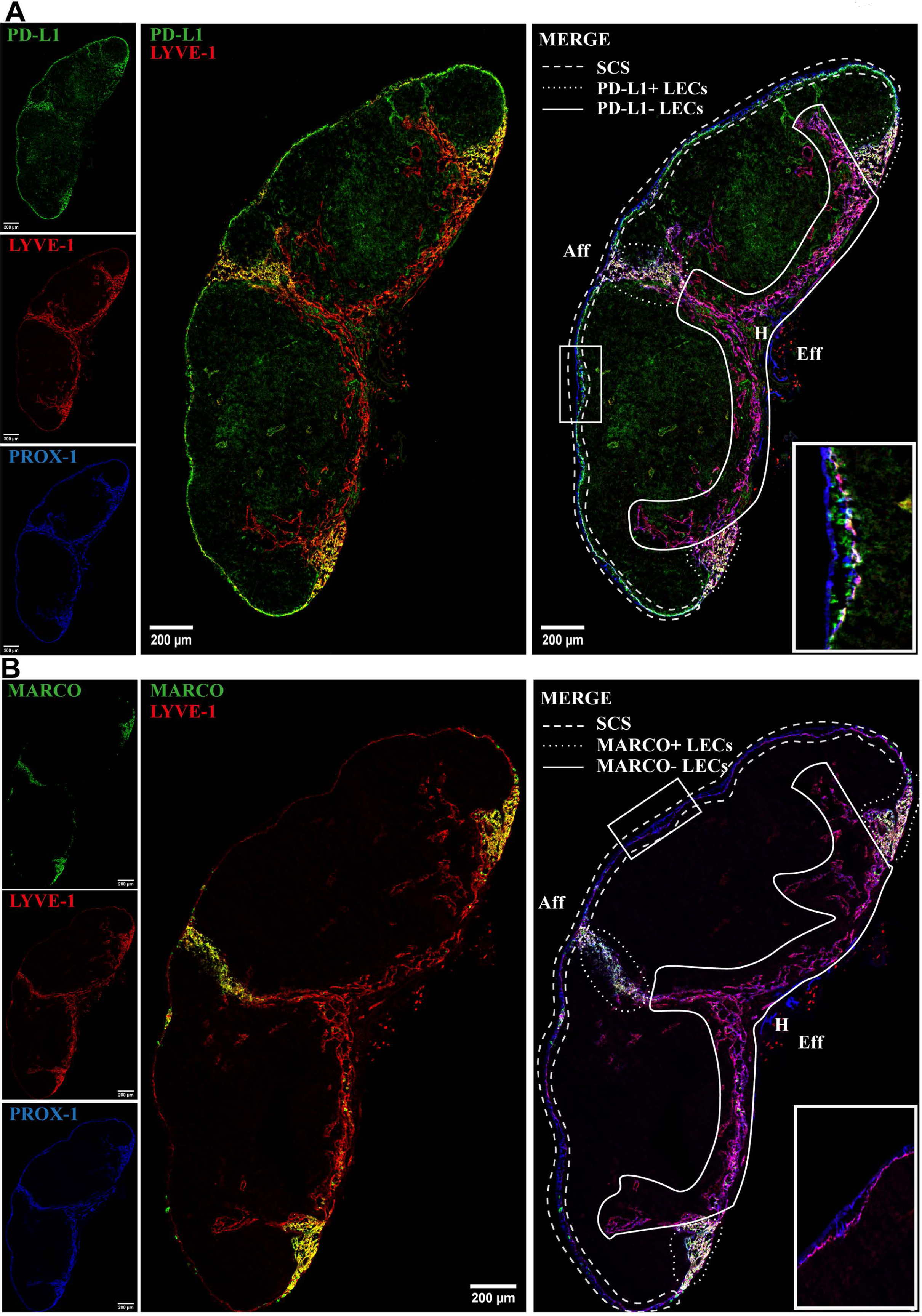
*In situ* mapping of lymph node LEC subsets. Serial sections of inguinal LN from *Prox-1-GFP* transgenic mouse. Immunoreactivity to PROX-1 (i.e. GFP) (blue), LYVE-1 (red) and either PD-L1 **(A)** or MARCO **(B)** (green). White lines indicate the different spatial location of lymphatic vascular niches: SCS LECs (dashed line); PD-L1+ **(A)** and MARCO+ **(B)** medullary sinuses (dotted line); PD-L1-**(A)** and MARCO-medullary sinuses **(B)** (solid line). The LN hilus (H), the afferent (Aff) and efferent (Eff) side of the LN are marked. Scale bar = 200µm. Insets show the SCS in higher magnification. Data are representative of three or more independent experiments.

As noted above, trajectory analysis predicts a close relationship between Marco-LECs and fLECs, and identifies a transitional population tzLECs between them. tzLECs are characterized by variable and often intermediate expression of fLEC (e.g. *Madcam1 and Ccl20*) and Marco-LEC associated genes (e.g. *Marco and Lyve1*) (Figure 1D). Histologic analysis identifies a region of LECs between fLECs and Marco-LECs that displays expression patterns consistent with this transitional phenotype. Cells of this phenotype can be observed between the fLECs and Marco-LECs in the region between the bilateral lobes of the inguinal LN and at the LN margins, where the lining cells display reduced MAdCAM-1 compared to the majority of fLEC and increasing levels of LYVE-1 (Figure 3). Few if any genes are specific to tzLECs (i.e. not shared with fLECs or Marco-LECs), and unbiased non-negative matrix factorization does not identify a gene set specific to the subset (data not shown). Thus, the tzLECs highlighted here are part of a continuum between fLEC and Marco-LEC populations, defining an area of vascular zonation. Consistent with computational predictions, the results favor a model in which Marco*-*LECs occupy the peri-follicular medulla in association to the LN cortex, with the transitional phenotype tzLECs bridging them to the SCS floor (Figure 4A).

**Figure 3:**
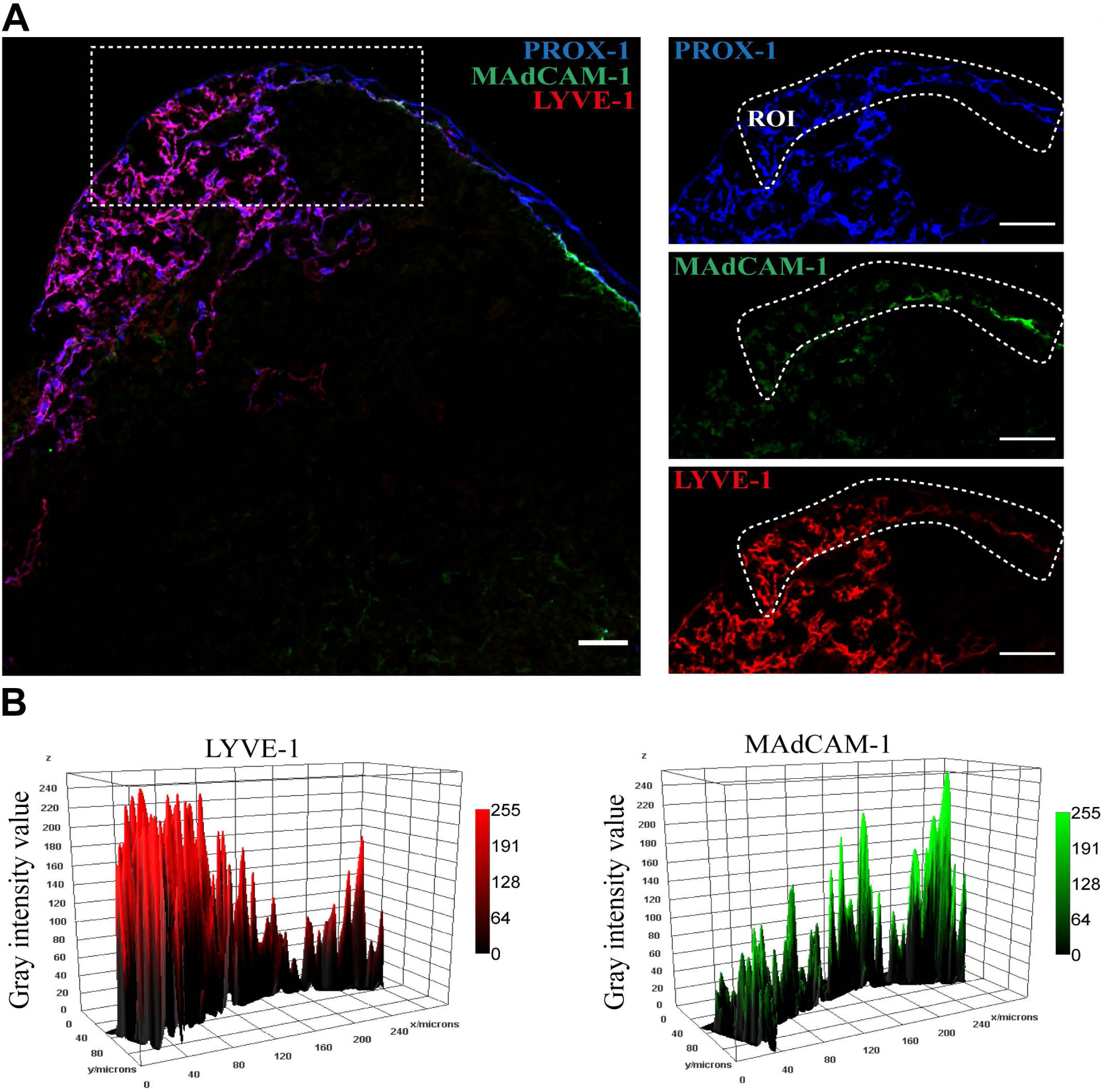
Transition zone (tz): LECs bridging fLECs and Marco-LECs. **(A)** Confocal microscopy of inguinal LN from *Prox-1-GFP* transgenic mouse. Immunoreactivity to PROX-1 (i.e. GFP) (blue), LYVE-1 (red) and MAdCAM-1 (green). Region of interest (ROI) is indicated. **(B)** Gray value intensity across the ROI for LYVE-1 (Marco-LEC marker) and MAdCAM-1 (fLEC marker) displaying decreasing or increasing expression along a path from the peri-follicular medullary sinuses to the SCS floor respectively. Scale bar = 50µm.

Ptx3-LECs are *Lyve-1* high but lack detectable *Pd-l1* (*Cd274*) and *Marco* expression (Figure 1D). This is consistent with the protein expression pattern of the central medullary sinuses in connection to the LN hilar region (Figure 2). These PD-L1-, MARCO-sinuses are associated with a network of scattered B lineage cells (Figure S2), a typical feature of the medullary region in the LN, where the lymphatic cords are surrounded by plasma cells [4; 34]. Trajectory analysis predicts that Ptx3-LECs are contiguous with the SCS ceiling (Figure 1C), which is also evident along the efferent side of the LN where the LYVE-1-outermost endothelial layers (cLECs) connect to the LYVE-1+ cords (i.e. Ptx3-LECs) (Figure S3). Hence, we have demonstrated the organization of LEC *in situ*, with SCS ceiling converging on Ptx3-LECs, Ptx3-LECs to Marco-LECs and tzLECs, then leading to fLECs (Figures 2 and 3), which mirrors the phenotypic progression revealed by trajectory analysis (Figure 1C). A schematic of the LN LEC connections is shown in Figure 4A.

### LEC molecular phenotypes correspond with their vascular niches and functions

#### The cLEC molecular profile reflects interaction with the LN capsule

cLECs connect to afferent and efferent collecting vessels, and co-express several genes with valve-related LECs, including higher levels of *Foxc2* [35] compared to other LN subsets (Figures 1D and S1C). Deletion of *Foxc2* in lymphatic vessels has been shown to lead to downregulation of the cLEC marker *Ackr4* but also to mislocation of surrounding smooth muscle cells [35]. cLECs interact physically with the relatively rigid capsule that surrounds LNs. The capsule is a dense network of connective tissue and hence is a unique extracellular matrix (ECM), distinct from other parts of the LN [4]. Reflecting this, cLECs are highly enriched for genes encoding ECM proteins, *Multimerin protein 1* and *2* (*Mmrn1* and *Mmrn2*), together with high levels of *Platelet-derived growth factor a* and *b* (*Pdgfa* and *Pdgfb*), *Jagged 1* (*Jag1*) (a ligand for Notch receptors on both endothelial and smooth muscle cells), and *Endothelin 1* (*Edn1*) (a vasoactive peptide and known lymphatic constrictor), proteins known to maintain interaction between endothelial cells and mural cells [36; 37] (Figures 4B and S4). The capsule was recently shown to have a CD34+ stromal cell subset with expression of CD248 [6], a ligand for MMRN2. MMRN2 can also interact with cLEC expressed MMRN2-ligand *Cd93*, which was recently coupled to ECM fibrillogenesis in tumor blood endothelium [38]. cLECs express the BMP family members *Growth differentiation factor 10* (*Gdf10*) and the *Bone morphogenic protein 4* (*Bmp4*) which differentiate cLECs from fLECs as well as bridging cells in the SCS (Figures 4B, S4 and S5). fLEC instead express *Bmp2* (Figures 4B, S4 and S5). The expression of BMPs may mediate both autocrine and paracrine signaling to the surrounding stromal and immune cells.

**Figure 4:**
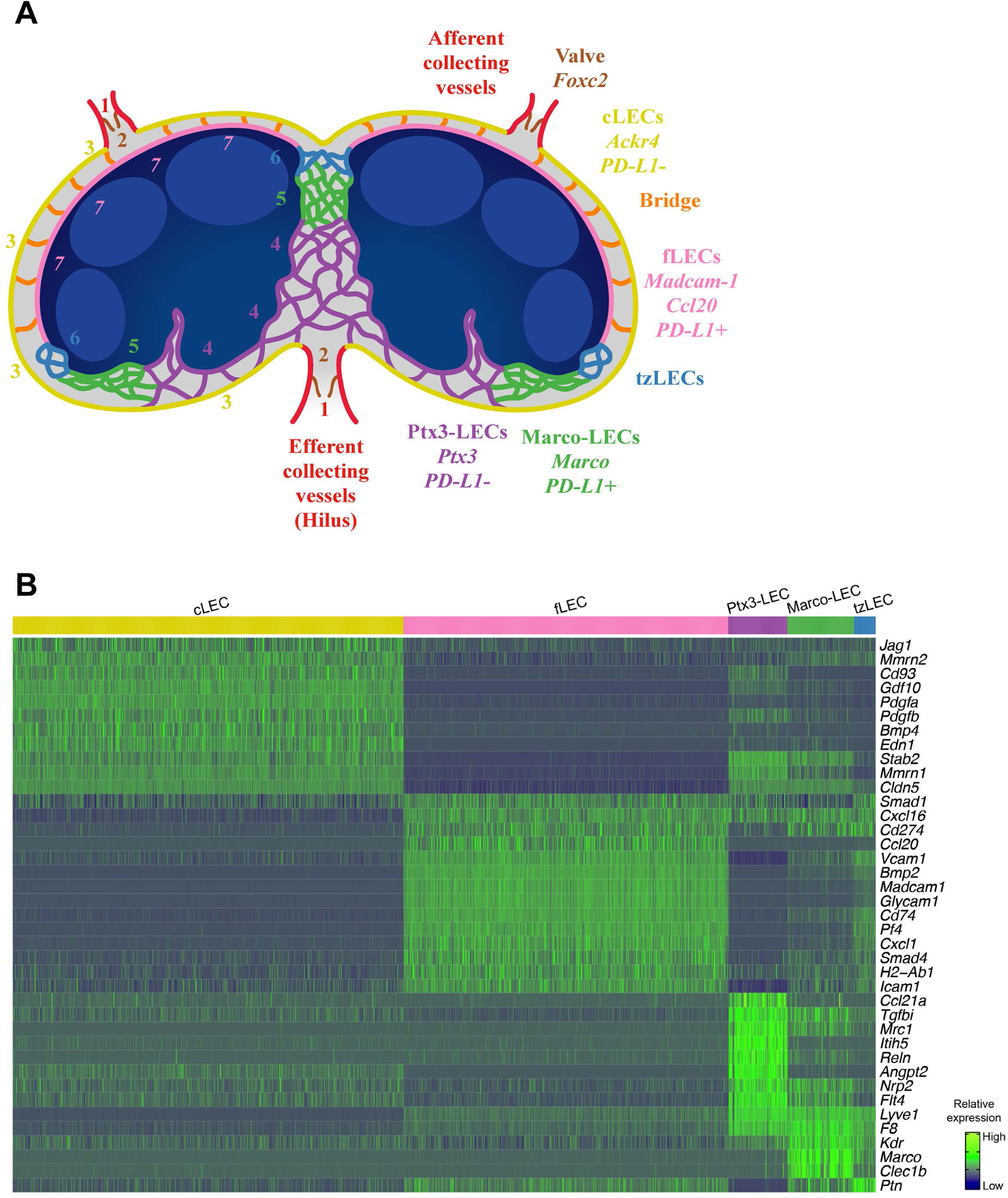
Mouse LN subsets and differently expressed genes (DEGs). **(A)** Cartoon showing the interconnections of LN LEC subsets related to tSpace alignments (Figure 1) and *in situ* mapping (Figure 2 and 3). **(B)** Heatmap of select DEGs in the C57BL/6 mouse (10x) LN LEC subsets. Values are imputed log counts (row scaled).

**Figure 5:**
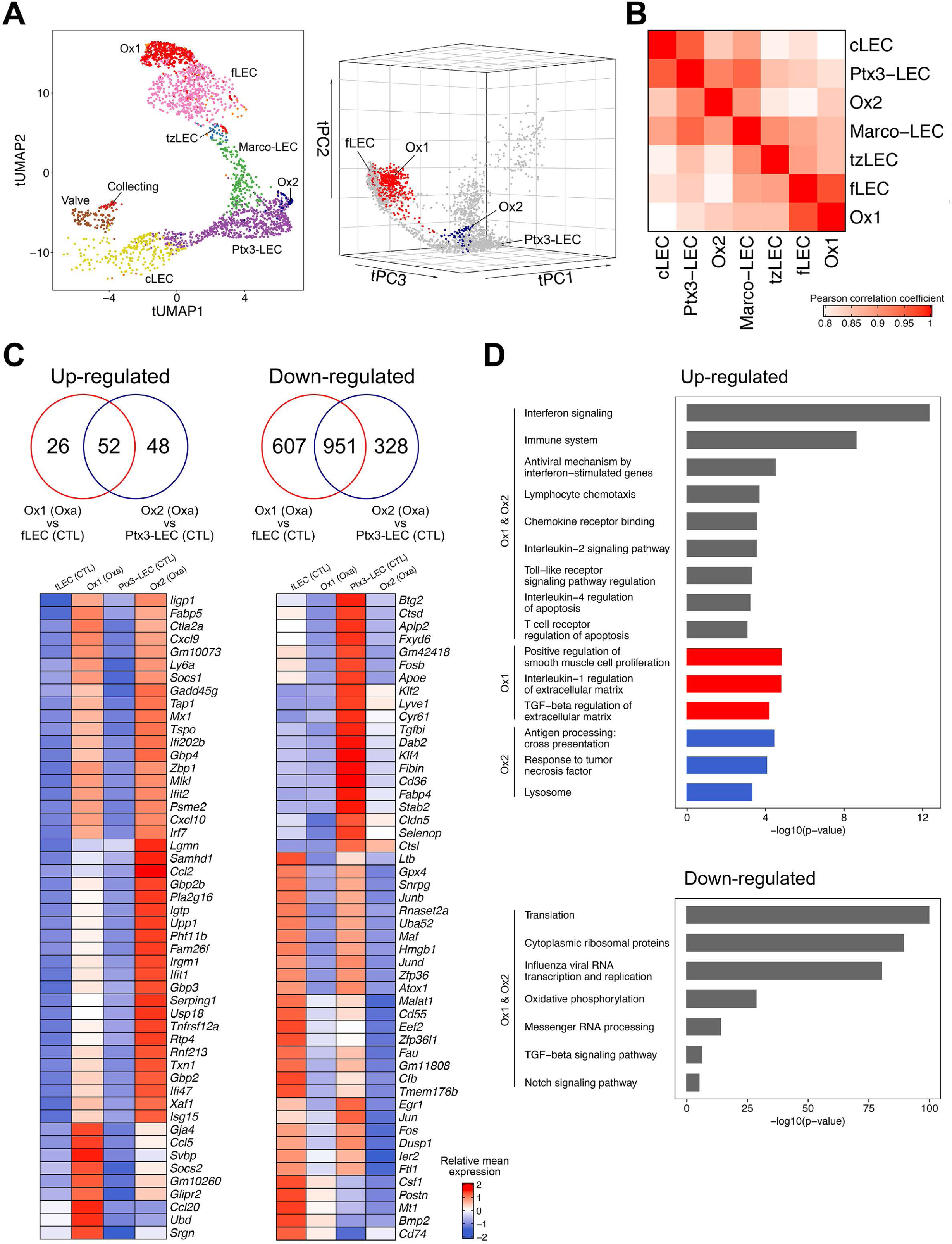
Niche-specific inflammatory responses in LN LECs. **(A)** tUMAP (UMAP on trajectory distances) of LEC from Oxa-treated and control mice, colored by subsets, including the two additional subsets (Ox1 and Ox2) in the Oxa-treated group (left). tPC (principal components on trajectory distances) of LEC from Oxa-treated and control mice, highlighting Ox1 and Ox2 respectively (right). **(B)** Pairwise Pearson correlation of LEC subsets in Oxa-treated group, calculated using the subset mean expression of the top 1000 variable genes. **(C)** Venn diagrams of DEGs from Ox1 in Oxa-treated LNs compared with fLEC in control LNs, and from Ox2 in Oxa-treated LNs compared with Ptx3-LEC in control LNs (upper; *p* < 0.01, fold change > 1.2). Heatmaps of 50 select upregulated or downregulated DEGs, with ribosomal genes excluded from the downregulated panel (lower; row scaled). **(D)** Select GO terms and BioPlanet pathways from Enrichr analysis of DEGs.

#### fLECs: an immune-active subset

fLECs are the gatekeeper for lymph-derived immune cell entry into the lymph node parenchyma. In keeping with this role, fLECs are characterized by genes involved in immune cell adhesion including *Madcam-1, Icam-1, Vcam-1* and *Glycam-1*, supporting active immune cell migration (Figures 1D, 4B and S4). These adhesion receptors may also help retain the closely associated SCS macrophages [16]. fLECs are also enriched for chemokines, including the known fLEC marker *Ccl20* (CCR6 ligand) [14; 26] but also *Cxcl1* (CXCR2 ligand) (Figures 1D, 4B and S4). CCL20 has been linked to Innate Lymphoid Cell (ILC) trafficking across the SCS [14] and may affect the cross-talk between CCR6 positive DCs and LECs in antigen presentation [39]. CCL20 could also influence B-cell homeostasis, as memory B-cells precursors are distinguished by high CCR6 expression in lymphoid organs of both mouse and man [40] and are closely associated to the SCS [41]. CXCL1 may instead contribute to neutrophil migration and recruitment [42]. fLECs and Marco-LECs, both macrophage-rich niches [16], share high expression of *Csf1* (Figure 7), and fLECs express *Platelet factor 4* (*Pf4*) (also known as CXCL4) (Figures 4B and S4). CSF-1 and CXCL4 both regulate monocyte and macrophage functions [17; 43] and LEC-derived CSF-1 is required to maintain the LN macrophage niches [17]. As mentioned earlier, *Bmp2* is specifically expressed by fLECs (Figures 4B, S4 and S5) that are also enriched for both *Smad1* and *Smad4* (Figures 4B and S4) suggesting a role for BMP-signaling in fLEC homeostasis.

**Figure 6:**
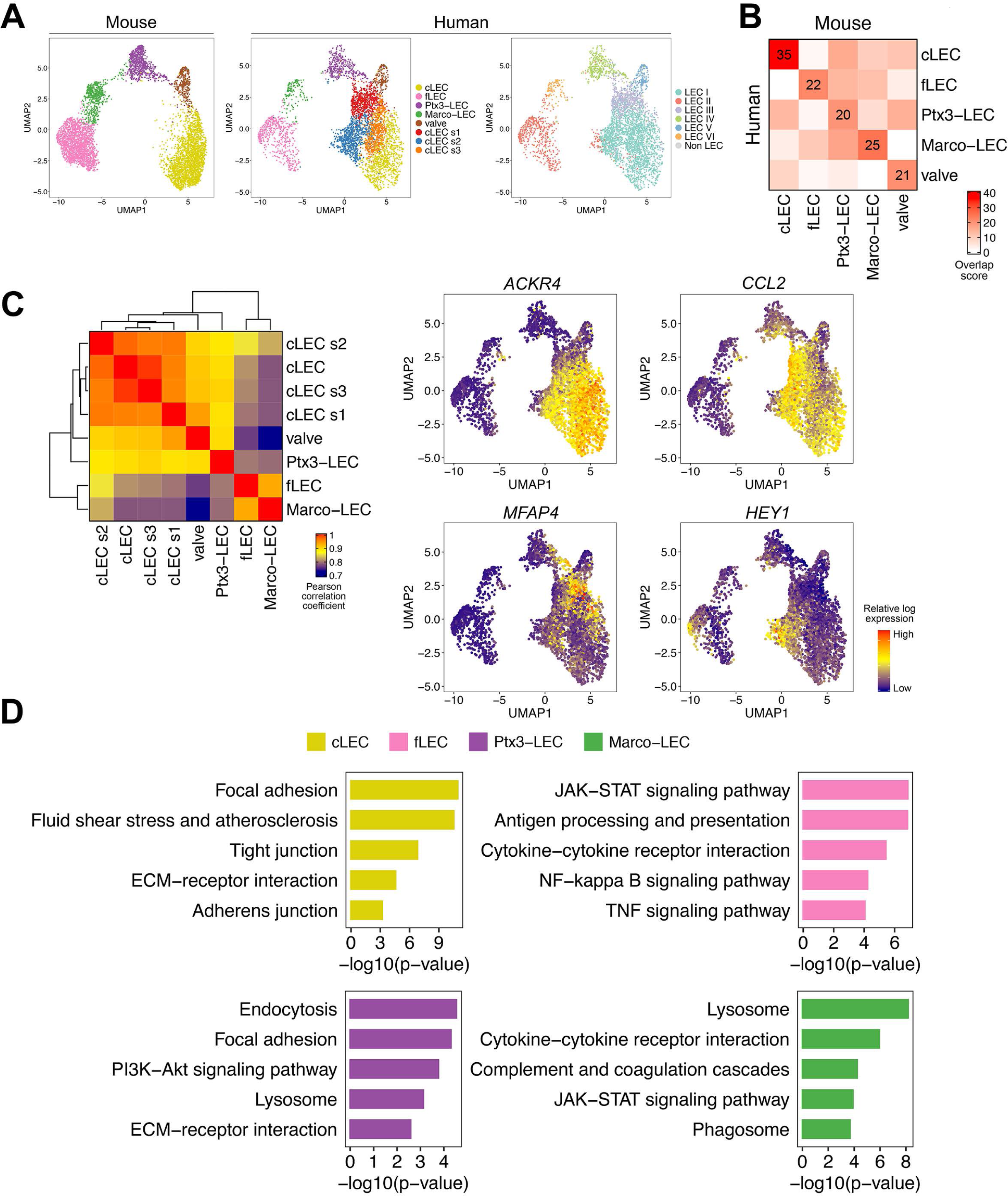
Cross-mapping of mouse and human LNs reveals conserved LEC populations and additional human cLEC subsets. **(A)** UMAP of aligned mouse and human LEC, colored by integrated mouse and human LEC subset (left and middle), or human LEC ID as in [26] (right). Bridge cells are not shown. **(B)** Pairwise overlap scores of top 100 subset-specific DEGs for mouse and human LEC. Overlap score is defined as the ratio between the number of shared genes observed and the number of genes expected to be shared by chance. **(C)** Correlation of gene profiles of human LEC subsets. Color scale: pairwise Pearson correlation coefficient, calculated using the mean expression of the top 1000 variable genes (left). Expression of cLEC subset-specific genes, projected on UMAP plot of human LN LECs (right). **(D)** Select pathways from Enrichr analysis of DEGs common to mouse and human.

**Figure 7:**
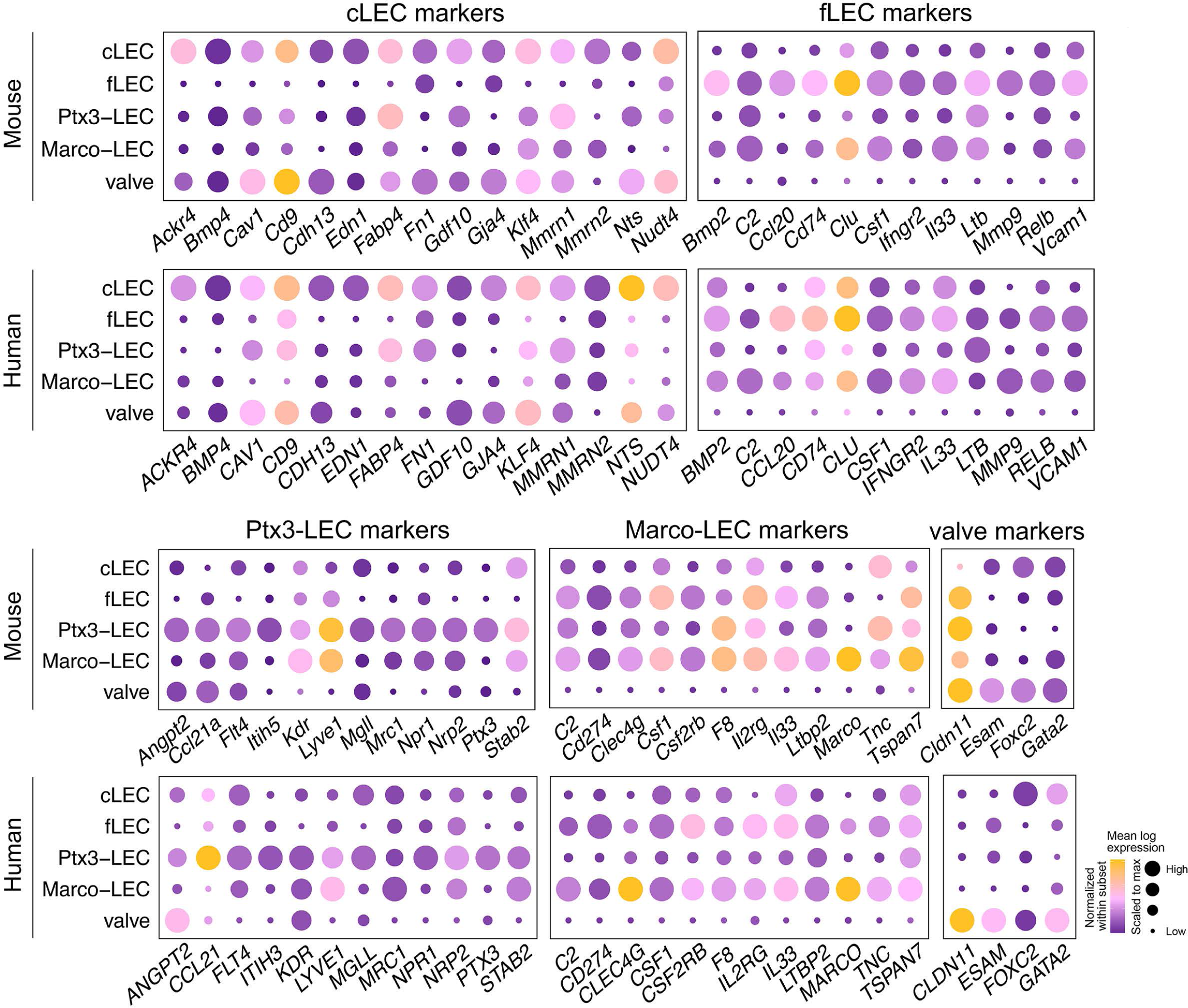
Conserved gene programs for mouse and human LEC specialization. Expression of select gene homologs (or paralogs *Itih5* and *ITIH3*) in cLECs, fLECs, Ptx3-LECs, Marco-LECs and valve LECs. Color indicates subset mean log transcript count, normalized within each subset. Dot size indicates subset mean log transcript count, scaled to the maximum value for each gene.

Another distinguishing feature of fLECs is their expression of *CD74* (Figures 4B and S4). CD74 is involved in the formation and transport of MHC class II-antigen complexes [44]. Together with high levels of *H2-Ab1* (Figures 4B and S4), this supports a major role of this subset in LN LEC-mediated antigen presentation. Together with the high expression of PD-L1 (*Cd274*) in fLECs (Figures 2, 4B and S4), and given that antigens are continuously transported from afferent lymphatics into the SCS, this places the fLECs as the major niche for LN LEC-mediated tolerance.

In contrast to all other subsets, fLECs and tzLECs lack expression of the scavenger receptor *Stabilin 2* (*Stab2*), a sinusoidal endothelial marker [45], and have low expression of the tight junction protein *Claudin-5* (*Cldn5*) (Figures 4B, S4 and S6). The latter suggests that junctional properties of the fLECs differ compared to other LN LECs and may reflect requirements for the active immune migration at this site.

#### Marco-LECs: Marco+ medullary sinus LECs

Immunofluorescence staining for MARCO in mouse inguinal LN shows that it delineates the medullary sinuses adjacent to the B cell follicles in the LN cortex (peri-follicular sinuses). MARCO expression coincides with PD-L1 expression in the medullary sinuses, and is excluded from the PD-L1 negative medullary sinuses closest to the hilus (Figures 2 and S2). Lymph from SCS passes through this lymphatic meshwork, abundant in macrophages that can capture lymph-borne pathogens [16; 46]. While MARCO is also marker for medullary macrophages [46], most of the MARCO expression within the adult LN is however confined to LECs (Figure 2). MARCO expression is also abundant in human LN medullary sinuses [26; 33]. In macrophages, MARCO expression facilitates phagocytic clearance by binding both gram-negative and positive bacteria [47], but its function in LECs is not known. Marco-LECs also express the C-type lectin *Clec1b* (CLEC2) (Figures 4B and S4), a ligand for Podoplanin (PDPN). CLEC2 is also expressed by myeloid cells, including DCs [48]; and interaction between PDPN+ fibroblastic reticular cells (FRCs) and CLEC2+ DCs has been implicated in FRC contractility and LN expansion in inflammation [48; 49]. LN LEC-expressed CLEC2 may mediate both homotypic and heterotypic cell interactions, since PDPN is highly expressed by LECs and by surrounding FRCs. *Stab2* is highly expressed by Marco-LECs and Ptx3-LECs (Figure S6), and together with Ptx3-LECs they are also enriched for the *coagulation Factor VIII* (*F8*) (Figures 4B and S4). We previously showed that lymph node LECs are a major source of FVIII production [50]. The present result shows that *F8* along with *Stab2* are features of medullary LEC subsets. Their co-expression may be a general feature of sinusoidal EC, since both are also characteristic of hepatic sinusoidal endothelium. FVIII forms a complex with von Willebrand factor (vWF) in plasma, a complex that is cleared by STAB2-expressing liver sinusoidal blood endothelial cells in a vWF-dependent manner [51]. However, the very low expression of vWF by mouse LN LECs [50] (and data not shown) suggests that FVIII may function as a pro-coagulant factor independent of vWF, possibly promoting the formation of fibrin mesh to block the spread of pathogens such as *Staphylococcus aureus* in the medulla [52].

The results show that Marco-LECs share gene expression patterns with myeloid cells, suggesting that this LEC subpopulation has a major role in innate immune functions. Interestingly, Marco-LECs and Ptx3-LECs express higher levels of *Kdr*, also known as *Vegfr2*, than other LN subsets (Figures 4B and S4), which is discussed in further detail below.

### Ptx3-LECs: *Ptx3+* central medulla and paracortical sinus LECs

*Marco*- and *Pd-l1-* (*Cd274*-) Ptx3-LECs uniquely express *Pentraxin-3* (*Ptx3*) (Figures 1D and 7). They are also distinguished by expression of *Inter-α-trypsin inhibitor 5* (*Itih5*), *Mannose receptor C-type* 1 (*Mrc1*)*, Reelin* (*Reln*) and a relative enrichment in *Ccl21a, Lyve-1* and *Stab2* (Figures 4B and S4). PTX3 belongs to the pentraxin family, secreted proteins with a cyclic and multimeric structure [53]. They bind pathogens and damaged self-proteins, acting as soluble pattern recognition molecules that mediate activation of complement and promote phagocytosis [53]. Medullary sinuses also support sustained and close interactions with long-lived plasma cells [34] and thus adaptive (antibody-based) and innate responses are likely to provide complementary defense mechanisms at this site.

Ptx3-LECs share gene expression with peripheral capillary LECs, with which they share morphologic features. Both have sprout-like blind ends specialized for attracting leukocytes and fluid flow into the lymphatics [4; 54] (Figure 2). *Ptx3* itself is not expressed by peripheral capillary LEC but capillary and Ptx3-LECs share *Itih5*, *Mrc1, Ccl21* and *Lyve-1* [55; 56; 57] (and data not shown). CCL21 mediates recruitment of CCR7+ dendritic cells (DCs) and T cells into peripheral lymphatic capillaries [57; 58]. Although in the peripheral LN Ptx3-LECs analyzed here *Ccl21* expression is relatively low (compare with levels in candidate collecting vessels in Figure 1D), it is more highly expressed in Ptx3-LEC in mesenteric LN (not shown) and in the human, as discussed below. Thus, CCL21 in Ptx3-LECs may contribute to leukocyte-LEC interactions and egress of CCR7+ cells including naïve B and T cells, possible complementing the known role of sphingosine-1-phosphate (S1P) in this process [54; 59]. Expression of the hyaluronan receptor Lyve-1 promotes the transmigration of DCs into blind-ended capillary lymphatic vessels [56]. MRC-1, a known marker for subsets of macrophages, binds microorganisms and promotes phagocytosis [47]. MRC-1 expression in peripheral lymphatic vessels also facilitates interaction with CD44-expressing immune cells [55], and could thus interact with activated CD44_high_ T-cells in egress from the LN. Thus, the paracortical and medullary sinus Ptx3-LECs share with peripheral capillary LECs gene programs for LEC-immune cell interactions, and support lymphocyte entry into medullary sinuses and exit from the LN.

*Ptx3* itself, as well as other genes selective to Ptx3-LECs, have known or proposed roles in the extracellular matrix (ECM). PTX3 binds collagen and fibrinogen-domain containing proteins, including ECM components, but also other pattern recognition molecules like C1q and ficolins [60]; thus it may amplify innate pathogen recognition mechanisms as well as contributing to maintenance and repair of the LN medullary environment. The Ptx3-LEC marker Reelin (*Reln*) is an extracellular matrix glycoprotein. In the periphery, Reelin is associated with the transition of peripheral capillary vessels to collecting vessels and contributes to the cross-talk between collecting vessel and surrounding smooth-muscle cells [61]. ITIH5 was originally isolated in a complex with hyaluronan [62] which can stabilize the ECM [63]. Notably, Ptx3-LECs also express high levels of the hyaluronan receptor LYVE-1 and ITIH-proteins have been shown to interact with PTX3 [64]. This provides a possible functional link between LYVE-1, ITIH5 and PTX3 in the Ptx3-LEC medullary sinus niche. Ptx3-LECs are also enriched for the collagen and integrin-binding ECM protein *TGFbeta-induced protein* (*Tgfbi*) (Figures 4B and S4), which is induced in LECs by hypoxia [65]. Together with high levels of the ECM protein *Mmrn1* (Figures 4B and S4), these genes support the notion of a highly specialized medullary ECM.

Ptx3-LECs show the highest levels of *Vascular endothelial growth factor receptor 3* (*Vegfr3*, or *Flt4*) [66] and of its co-receptor *Neuropilin 2* (*Nrp2*) [67] (Figures 4B and S4), suggesting a higher responsiveness to Vascular Endothelial Growth Factor C (VEGFC) [68; 69]. VEGFC is a major driver of lymphangiogenesis in development [68] and in the adult lymphatic vasculature [70]. VEGFR2 (*Kdr*), which binds both VEGFC and VEGFA [66], is also highly expressed in Ptx3-LECs, although it is most enriched in Marco-LECs. Thus, VEGFA and VEGFC may elicit niche-specific responses in medullary sinus LECs, acting differently on *Vegfr3*_high_ *Vegfr2*_intermed_. Ptx3-LECs and *Vegfr3*_intermed/*low*_ *Vegfr2*_*high*_ Marco-LECs. Interestingly, while SCS fLEC and cLECs are established in embryogenesis [35], the medullary sinuses form by sprouting during the first two postnatal weeks [71]. Thus high *Vegfr3* (*Flt4*) in Ptx3-LEC could reflect retention of their postnatal programs for VEGFC-dependent sprouting [71], and may imbue them with the capacity to regenerate or expand rapidly in response to the requirements of LN inflammatory responses. The Ptx3-LEC marker *Angiopoietin 2* (*Angpt2*) has also been linked to LEC proliferation [72; 73]: it can induce lymphatic hyperplasia when overexpressed [73] and is highly expressed by newly formed lymphatic vessels in inflammation-induced lymphangiogenesis [72]. Ptx3-LECs, like cLECs, but in contrast to all other parenchymal LN LEC subsets, also lack expression of PD-L1 (*Cd274*). Loss of PD-L1 is associated with increased LN LEC proliferation but also increased LEC apoptosis after inflammation [24]. Thus, low PD-L1 suggest higher responsiveness not only to proliferation-inducing signals but also higher sensitivity to apoptotic cell death, the latter thought to facilitate LN contraction after inflammatory stimuli [24]. Taken together, high *Vegfr3, Vegfr2, Nrp2, Angpt2* and low *CD274* (PD-L1) suggest a major role of Ptx3-LECs in LN remodeling.

### A cellular bridge between the SCS ceiling and floor

As discussed earlier, trajectory analysis not only predicts a transitional population, tzLECs, that link fLEC and Marco-LEC, but further identifies a bridge population that connects cLEC and fLEC. This bridge population is variably represented in the mouse samples (Figures 1B and S1A) but is prominent in the human (see below, Figure 9). We plotted gene expression of cells along the path from cLEC to fLEC (Figures S5 and S7A) and found that individual cells within the bridge display variable levels of cLEC and fLEC marker transcripts including *Lyve1* (Figure S7A), but overall a decreasing gradient of cLEC markers (e.g. *Ackr4* and *Cav1*) and an increasing gradient of fLEC markers (*Madcam-1*, *Pd-l1/Cd274*) in the progression from cLEC to fLEC (Figure S7A). We speculate that these cells participate in formation of previously described “transendothelial channels”, physical bridges that link the subcapsular ceiling and floor [10]. Immunofluorescence imaging indicate that these cords consist not only of channels [10; 35], but also nucleated LECs traversing the sinus, as indicated by nuclear staining for the lymphatic reporter *Prox-1-GFP* (Figure S7B). We used immunofluorescence and *in situ* hybridization for RNA (RNAscope) to evaluate bridge cell expression of cLEC and fLEC markers. Most bridging cells displayed defining phenotypes of cLEC (LYVE-1-, PD-L1-, MAdCAM-1-) or fLEC (LYVE-1+, PD-L1+, MAdCAM-1+) on the protein level: cells co-expressing these cLEC and fLEC markers (as predicted for cells of the computationally defined bridge) were rare or absent (Figure S7). However, RNAscope revealed that bridging cells lack mRNA for *Bmp4* (cLEC marker) and have intermediate expression of *Bmp2* (fLEC marker), a pattern consistent with the single cell profiles of the predicted bridge (Figure S5). Although this observation provides some support for the correspondence between the physical and trajectory-inferred cellular bridges, they are far from being conclusive. An alternative possibility is that fLEC and cLEC can interconvert in response to local environmental signals, and that the transcription profiles of these “bridge” cells are snapshots of cells in process of these transitions. Further experiments are required to map the computationally defined “bridge” cells within the LN lymphatic network, and to confidently identify the profiles of the *in situ* cellular links between SCS floor and ceiling.

**Figure 8:**
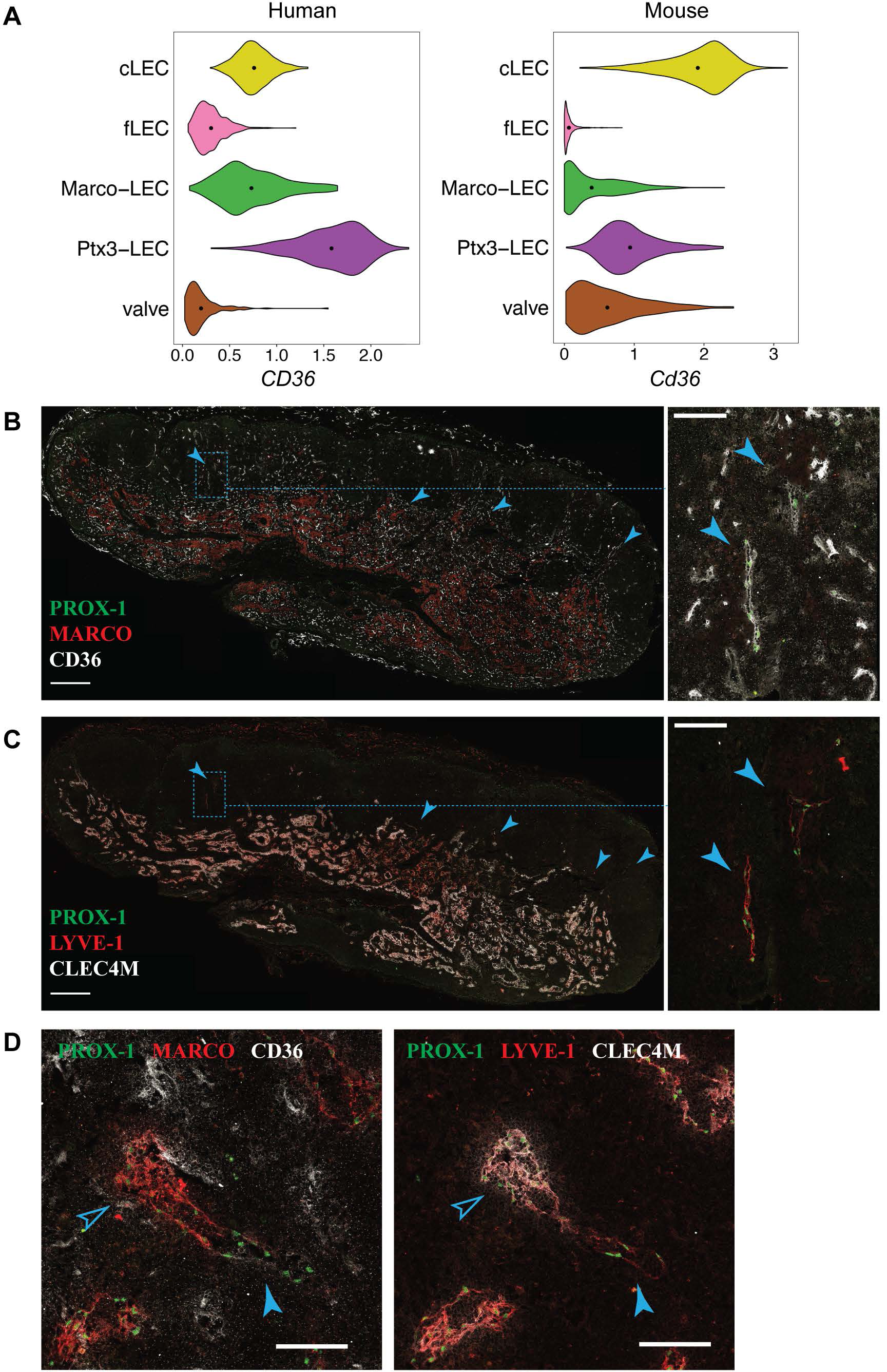
*In situ* localization of Ptx3-LECs and transition between Ptx3-LECs and Marco-LECs in human LNs. **(A)** Expression of *CD36/Cd36* in LN LEC subsets of human and mouse. Dots indicate mean log-normalized transcript count. **(B-D)** Identification of CD36_high_ Ptx3-LECs in human head and neck LNs by immunostaining. **(B, C)** Immunofluorescence of PROX-1, MARCO and CD36 **(B)**, or PROX-1, LYVE-1 and CLEC4M **(C)**. Zoomed-in images (inset marked by blue dotted lines) in B and C demonstrate CD36_high_ LYVE-1+ paracortical sinuses (filled arrowhead). Scale bars = 500 µm (left panels) and 100 µm (right panel inset). **(D)** CD36_high_ LYVE1+ Ptx3-LECs (filled arrowhead) can be seen associated with MARCO+ CLEC4M+ Marco-LECs (empty arrowhead) in human LNs. Scale bars = 100 µm. CD36_high_ Ptx3-LECs were detected in four out of seven human LNs. Images are representative of four biological replicates.

**Figure 9:**
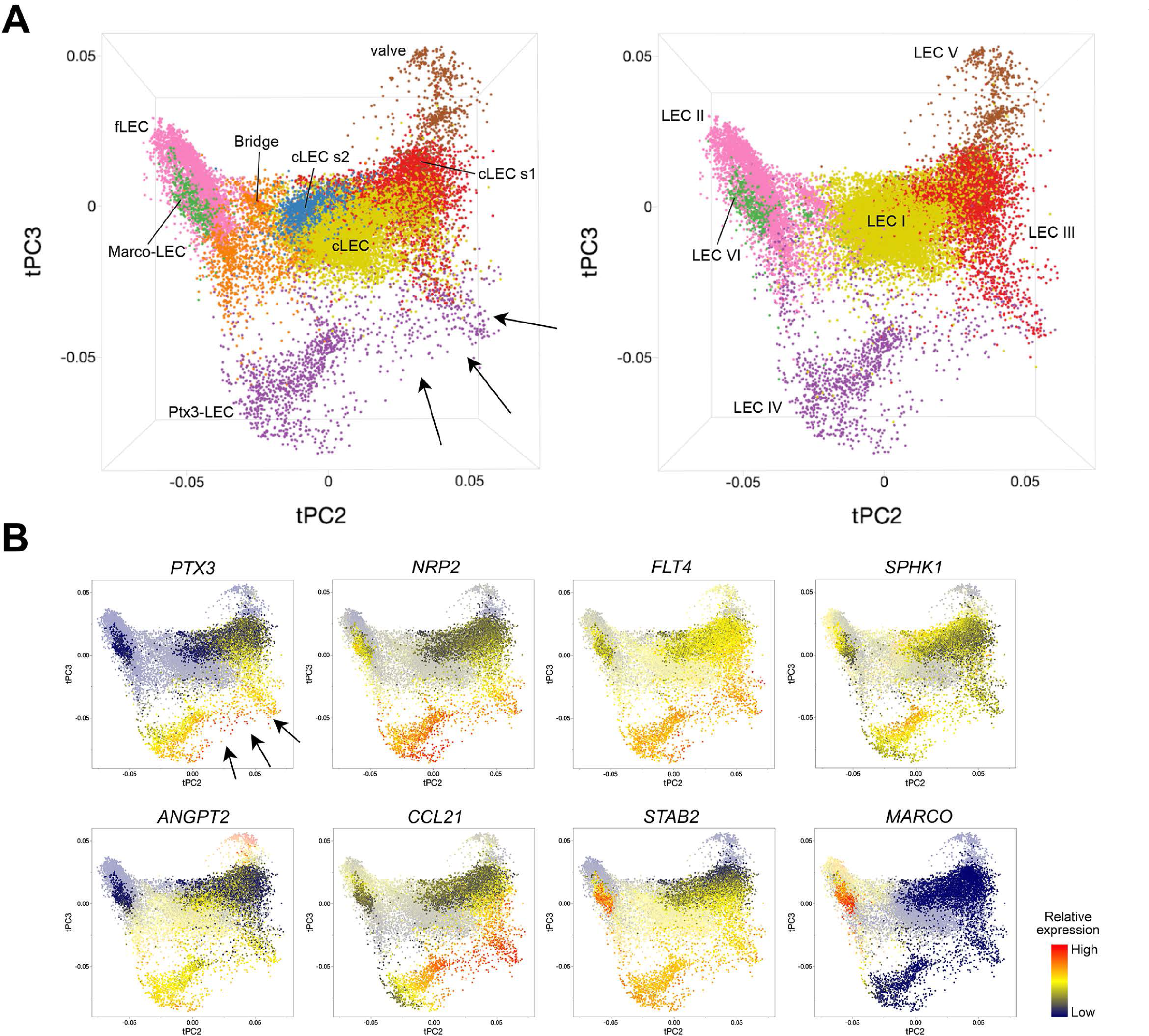
A continuum of human LEC phenotypes in trajectory space. Principal component projections of trajectory distances reveal complex zonation of LEC subsets and intermediates. **(A)** 3D projections with LEC subset identities defined by correlation with populations in the index human sample (left) or by unsupervised clustering as described in [26] (right). **(B)** Expression pattern of indicated genes. Marco-LEC, Ptx3-LEC, and cLEC s1 populations are highlighted. Values are batch-corrected imputed log counts.

### Niche-specific inflammatory responses in LN LECs

Oxazolone (Oxa) is a chemical compound used in a prototypical model for inflammation-induced LN hypertrophy [74]. We here chose to look at initial LEC responses 48 hours after topical skin application of Oxa, before any major lymphangiogenic responses are expected. Lymphangiogenesis is a late effect in LN remodeling, peaking at 5-7 days after Oxa, or other inflammatory stimuli [24; 74; 75]. We aligned LEC of the Oxa-treated mouse with the untreated control of the same genetic background and observed two new subsets, termed Ox1 and Ox2, in the immunized group (Figure 5A).

Inspection of the tSpace projections tUMAP (UMAP on trajectory distances) and tPC (principal components of trajectory distances) suggests that Ox1 arises from fLECs, and Ox2 from Ptx3-LECs (Figure 5A). Pairwise Pearson correlation based on the top 1000 variable genes confirmed that Ox1 is most correlated with fLECs (Pearson correlation coefficient = 0.97) and Ox2 with Ptx3-LECs (Pearson correlation coefficient = 0.92) (Figure 5B). We asked which genes were differentially expressed in Ox1 and Ox2 due to Oxa treatment (Figure 5C). Compared to fLEC and Ptx3-LEC in untreated LN respectively, we found that both Ox1 and Ox2 displayed typical interferon (IFN) responses, with dramatic induction of *Cxcl9* and *Cxcl10* as well as *Irf7*, the master regulator of type-I IFN-dependent immune responses [76] (Figure 5C). CXCL9 and CXCL10 bind CXCR3 which mediates recruitment of dendritic cells, NK cells, effector T-cells [77] and, in inflamed LNs, subsets of monocytes [78]. Both Ox1 and Ox2 also showed upregulated *Psme2*, a proteosomal component involved in peptide processing for class I antigen presentation [79] (Figure 5C).

While most gene changes were shared, some were preferentially or more dramatically increased in one or the other subset (Figure 5C). *Ccl20*, which is selectively expressed by fLEC in resting LNs (Figure 1D), is highly upregulated in Ox1 but not Ox2, maintaining its selective pattern of expression in the SCS floor (Figures 5C, 1D and 4B). Ox1 also displays selective upregulation of *Ubiquitin D* (*Ubd*), important in activation of the transcription factor NF-kappa B [80]. Oxa controls subset-specific upregulation of monocyte chemoattractants *Ccl2* (Ox2) and *Ccl5* (Ox1). CCL5 acts through CCR5, which is expressed by multiple adaptive and innate immune subsets; while CCL2 acts through CCR2 in monocyte recruitment [42]. Thus, chemotactic regulations in the SCS and medulla diverge. Gene enrichment analysis confirms Ox1 and Ox2 share enrichment in interferon signaling and immune system signatures, but the subsets also display unique ontologies: e.g. Transforming Growth Factor (TGF)-beta regulation of ECM in Ox1 and response to Tumor Necrosis Factor (TNF) in Ox2 (Figure 5D). Several of the observed changes, including induction of *Ccl2, Ccl5, Ccl20, Cxcl9* and *Cxcl10* were observed previously in analyses of bulk LN LECs in response to HSV-1 virus [75] or after ovalbumin specific T-cell responses [11]. Our results link inflammation-induced transcriptional changes to specific LEC subsets, implying that niche-specific changes coordinate LEC-driven responses in early inflammation

Interestingly, in both Ox1 and Ox2, significantly more genes are downregulated than upregulated (Figure 5C). Pathway analysis shows downregulation of genes associated with viral gene expression, ribosomes and mRNA processing (Figure 5D). This may be a result of the activation of type I interferon signaling (Figure 5D), which can cause widespread downregulation of host and viral transcriptional and translational pathways as a defense against viral replication [81; 82]. Downregulated genes include *Csf1* and *Ltb* (Figure 5C), which is interesting from the perspective of the known transient loss of sinusoidal LN macrophages in acute inflammation [17; 18]. Core lymphatic endothelial differentiation genes are also downregulated, including *Lyve-1 and Cldn5*; and subset marker genes are inhibited as well: Ox1 shows reduced expression of fLEC marker genes *CD74* and *Bmp2*; and Ox2 of Ptx3-LEC marker genes including *Stab2* and *Tgfbi* (Figure 5D). In addition to IFN-induced transcriptional repression, another contributing factor may be transcription factor availability, which can dictate level of gene expression [83]: TNF-induced RELA-dependent pathways, which are also induced as part of the Oxa response (Figure 5D), have been shown to redirect cofactors from super-enhancers, repressing cell identity genes in a cell type dependent manner [84].

### A conserved pattern of LEC phenotypes in human and mouse LNs

#### Mouse and human common LN LEC phenotypes

We have previously reported LEC diversity and described multiple LEC subsets in human LNs [26]. Here we compare our mouse LEC samples to a representative sample of human head and neck LN LECs, selected for its quality, high cell number and inclusion of each of the subsets we identified. We translated human gene names to their mouse homologs, and integrated human and mouse LECs using Canonical Correlation Analysis (CCA) [85], which implements a variant of the mutual nearest neighbors algorithm (MNN) [86]. Cross-mapping successfully aligned human and mouse LECs, and unsupervised clustering of the aligned cells identified shared but also unique subsets in human LN (Figure 6A). The five subsets common to mouse and human (cLEC, fLEC, Ptx3-LEC, valve-LEC and Marco-LEC) display high correspondence with human LEC subsets defined previously using unsupervised clustering [26] (LEC I, II, IV, V and VI respectively) (Figure 6A). Specifically, the Marco-LEC subset of medullary EC maps to human LEC VI, which we previously characterized as medullary sinus *CD209*+ LECs [26]. However, mouse Ptx3-LECs map to human LEC IV, which we postulated to be peripheral capillary LECs [26]. We discuss this below.

#### Specialization of the human SCS ceiling

Unsupervised clustering aligned the mouse cLECs with one subset of the human cLECs, “(core) cLECs”, which we identified previously as LEC I [26] (Figure 6A), but additional cLEC subsets (cLEC s1, s2 and s3) were unique to the analyzed human LN. Pearson correlation analysis reveals similarity of these subsets with the core cLECs (Figure 6C). cLEC s3 is most similar to the core cLEC population as indicated by Pearson correlation (Figure 6C), UMAP (Figure 6A) and differentially expressed genes (DEGs) (data not shown): we do not discuss this subset further. cLEC s1 corresponds to MFAP4+ SCS LEC (LEC III), which we located overlying the human LN medulla [26]. Consistent with this, cLEC s1 clusters close to Ptx3-LECs (Figure 6A), and links other cLECs to Ptx3-LECs in trajectory space (see below Figure 9A). cLEC s2, which we did not segregate previously, uniquely expresses high levels of *Hairy/enhancer-of-split related with YRPW motif protein* (*HEY1*), the chemokine *CCL2* (Figure 6C) and *E-Selectin* (*SELE*) (data not shown). Based on *HEY1* and *SELE* expression, the subset is identifiable in 3 of the 6 human LN samples we studied previously [26] (data not shown). The findings suggest greater heterogeneity and specialization in SCS ceiling LEC in humans than in the resting specific pathogen free (SPF) mice studied here. The subset specialization of human cLEC may relate to local differences in immune environments, or to the more complex architecture of the human LN. Unlike the mouse, human cLECs participate in invaginations of the capsule, known as trabecular sinuses [4], which may experience more turbulent flow of the incoming lymph and hence variation in shear stress.

#### Comparisons of human and mouse LEC differentially expressed genes (DEG)

To evaluate similarities between species, we compared DEGs of mouse and human LEC subsets using a gene overlap score, defined as the ratio of the number of shared DEGs to the number of genes expected to be shared based on random chance, for each combination of mouse and human subsets. In all instances, the highest overlap scores are seen between corresponding subsets (Figure 6B). Based on overlap scores, cLECs are more conserved than the more immunologically active subsets fLECs, Marco-LECs and Ptx3-LECs. That floor and medullary sinus subsets showed less conserved DEG profiles, likely reflects evolutionary pressure in response to pathogens, contrasting with conservation of structural functions of cLECs.

Gene set enrichment analysis based on conserved genes has the potential to highlight core functions of the LEC subsets (Figure 6D). Shared cLEC genes are involved in focal adhesion, fluid shear stress response and ECM interaction, consistent with their structural role and association to the capsule. Ptx3-LECs are enriched in genes for endocytosis and lysosome, which could relate to the high expression of scavenger receptors (e.g. *Lyve-1, Stab1, Stab2*). Marco-LECs are enriched for lysosomal genes, and for genes of the JAK-STAT signaling pathway as well as the coagulation and complement cascades. fLECs display high enrichment for inflammatory pathways, including JAK-STAT and Nuclear Factor (NF)-kappa B and are enriched for genes involved in antigen processing and presentation, supporting their immunomodulatory properties. A number of mouse key subset specific marker DEGs are shared in the human including e.g. fLECs: *Bmp2, Ccl20, Cd74 and Csf1*; cLECs: *Ackr4, Bmp4 and Cav1;* Marco-LECs: *Marco, Clec4g and C2;* and Ptx3-LECs: *Ptx3, Nrp2 and Flt4* (Vegfr3). We illustrate expression of these and other select shared marker genes in Figure 7.

As noted above, mouse Marco-LECs align with LEC VI, which we identified previously as the major medullary sinus subset in human LNs [26]. Unexpectedly, mouse Ptx3-LECs align with a subset (LEC IV) that we previously related to capillary LECs based on their enrichment for expression of *Ccl21* and *Lyve-1* [26], gene markers of peripheral capillary lymphatic vessels [56; 57]. However, these genes are also expressed by mouse Ptx3-LECs (Figures 1D, 4B, and 7), and as noted earlier, their expression likely reflects the parallels with capillary LEC in morphology (blind ends, loose EC junctions) and function (recruiting fluid and lymphocytes into lymph) [4]. Supporting a medullary identity, Ptx3-LECs and LEC-IV share high levels the sinusoidal endothelial marker *Stab2/STAB2* [45], shown to be expressed by medullary sinuses in human LNs [87]. Importantly, human Ptx3-LECs (LEC IV) also share expression of *PTX3* and lack *PD-L1* (*CD274*) expression, similar to their mouse counterpart; and they express the inter-alpha-trypsin inhibitor gene family member *ITIH3*, functionally related to the mouse Ptx3-LEC marker *Itih5* [63] (Figure 7).

Ptx3-LECs in both humans and mice are distinguished from Marco-LECs by higher expression of the glycoprotein and scavenger receptor *CD36,* also known as *Fatty Acid Translocase (FAT)* (Figure 8A). Staining of CD36 in human head and neck LNs identified capillary-like, CD36_high_ Lyve-1+ lymphatic cords, negative for MARCO and the Marco-LEC (LEC VI [26]) marker CLEC4M (Figures 8B and 8C). These CD36_high_ lymphatic sinuses were found either as isolated cords in the paracortex (Figures 8B and 8C) or as extended sprouts from MARCO+ CLEC4M+ medullary sinuses (Figure 8D), similar to mouse Ptx3-LECs which connects to Marco-LECs in perifollicular regions (Figure 2). Thus cross-species comparison of single-cell profiles (Figure 6) and *in situ* analysis (Figure 8) support that human and mouse share two distinct niches of paracortical and medullary sinus LECs: *MARCO+* LECs, which correspond to the previously published *CD209*+ medullary sinus LEC subset (LEC VI) [26] and Ptx3-LECs, CD36_high_ paracortical and medullary sinus LECs, corresponding to LEC IV [26] in our earlier classification of human LN LECs. Notably, human and mouse LN LECs also share the *in situ* physical transition between Marco-LECs and Ptx3-LECs, as predicted in trajectory analysis (Figures 1C and 9A, discussed below).

## Species-specific gene expression

A number of genes with homologs in mouse and human are not conserved in expression, or display different patterns of subset selectivity. Here we highlight select examples for discussion (Figure S8). As noted earlier, the mouse fLEC markers *Madcam-1* and *Msr1* are poorly expressed by human LECs [26] (data not shown). *ACKR1*, also known as *DARC* (Duffy Antigen Receptor for Chemokines) marks human fLECs and Marco-LECs; but *Ackr1* has very low expression in mouse LN LECs, without clear subset selectivity (Figure S8B). Consistent with this, endothelial ACKR1 in mouse is restricted to venular blood vessels, with only sparse and low detection in LECs [88]. ACKR1 is a chemokine “interceptor”, which can serve as a sink for a large range of inflammatory chemokines [89]: its expression could reflect a greater need to moderate inflammatory chemokines in the human. Alternatively, it may facilitate transport of chemokines across the lymphatic endothelium in the human LN, as ACKR1 has been shown to shuttle chemokines across endothelial cell layers [90]. Several ACKR1 ligands are expressed by human fLECs including CXCL3 and CXCL5 [26].

Human but not mouse LN LECs also display high expression of *IL-6* in fLECs (Figure S8B), likely reflecting a higher inflammatory basal state in human, especially compared to our SPF mice. Human Ptx3-LECs express *MMP2* and *LOX*, genes missing in mouse LECs or expressed in different subsets (i.e. *Lox* expressed weakly in cLECs in mouse) (Figure S8B). Both Matrix metalloproteinase 2 (MMP2) and Lysyl oxidase (LOX) can contribute to ECM modulation [91; 92], suggesting that, although matrix interplay is conserved, the specific mechanisms of matrix interaction in this population of LECs may have diverged. Mouse-specific LN LEC DEGs include *Apolipoprotein E* (*ApoE*) in cLECs and medullary sinus populations and *Regulator of G-protein signaling 4* (*Rgs4*) in fLECs (Figure S8B). *Carboxypeptidase E* (Cpe) and *Carbonic anhydrase 8* (*Car8*) are examples of genes with different expression pattern across mouse and mouse LN LEC subsets (Figure S8B). Since scRNA-seq is often unable to detect low abundance transcripts, the apparent lack of expression of a gene must be interpreted with caution: the expression pattern of the genes mentioned here have been observed in each of our samples.

In addition, CD209 and CLEC4M, which lack orthologues in mouse [26], are specific for human Marco-LECs. We described them previously as medullary sinus markers [26]. Human IL-32 and mouse Glycam1 also lack orthologues. IL-32 potently amplifies inflammatory responses by induction of multiple cytokines [93]; it is highly expressed by human Ptx3-LECs (data not shown). *Glycam-1,* a secreted mucin that on high endothelial venules is decorated with glycotopes for leukocyte selectins [94], is selectively expressed by mouse fLEC (Figures 4B and S4).

## The continuum of human LEC phenotypes in trajectory space

We have focused to this point on comparing “subsets” of human and mouse LECs, but as illustrated for the mouse, LECs exist in a phenotypic continuum that may reflect physical alignments (spatial transitions), developmental sequences, or both. To gain further insight into LEC diversity and zonation within human LNs we ran tSpace on combined LN LECs from our six previously published human samples [26]. In the tSpace projections in Figure 9A, cell identities are determined based on correlation with the core subsets in our index human LN (left) or based on unsupervised clustering as previously [26] (right). Alignment of human LECs in trajectory space reveals both similarities and differences to the mouse. Shared subsets are aligned in the same order as in the mouse, with links from Valve-LECs to cLECs, Ptx3-LECs, Marco-LECs and finally fLECs (Figure 9A) (Compare with mouse alignments in Figures 1B and 1C). However, in the human the cellular “bridge” between cLECs and fLECs is highly populated with cells. The human-specific cLEC subsets also show interesting alignments. cLEC s2 appears closely linked to the bridge. cLEC s1, which we previously identified as SCS ceiling LEC overlying the medulla (LEC III) [26], extends towards and links prominently to Ptx3-LECs (see arrows): Cells from both “subsets” within this Ptx3-LECs-to-cLEC s1 trajectory express *PTX3* itself, and are highly enriched in *CCL21* (Figure 9B), suggesting a relationship of *PTX3*+ medullary sinuses and the peri-hilar ceiling. Co-expression of *CCL21* and *SPHK1* (enzyme required for S1P production) support a role of Ptx3-LECs in lymphocyte egress. Heterogeneity in *ANGPT2* also exists along the Ptx3-LEC paths (Figure 9B).

The alignments outlined here are not an artifact of the high cell number or integration of LEC from different LNs, because the same patterns and linkages are seen in independent tSpace projections of our index LN sample (not shown). Human LN LECs thus display numerous intermediate phenotypes and complex zonation that may reflect the complexity of human LN architecture observed histologically.

## Summary and outlook

The LN lymphatic vasculature provides a complex lymphatic vascular bed, adapted to a unique microenvironment where cooperation between stromal cells and immune cells forms the premise for effective immune cell interaction and activation. Endothelial cells in this environment not only need to provide a framework for the structural organization of the organ, but also need to be able to adapt to constant changes induced by immunological stimuli, organ expansion in LN hypertrophy and to support tolerogenic immune reactions in homeostasis.

We define five LN LEC subsets in mouse: valve LECs, the SCS *Ackr4*+ cLECs, the immune active *Ccl20*+ fLECs, as well as Ptx3-LECs and Marco-LECs, two medullary sinus subsets that were not recognized previously. Single-cell gene profiles indicate niche-specific functional specialization of these subsets with distinct pathways for pathogen interactions and matrix modeling. Interestingly, cross-species mapping with human LN LECs shows that both subsets as well as their respective functions are conserved, which allows us to redefine the subset previously identified as candidate capillary LECs (LEC IV) [26] in human LN to paracortical and medullary sinus Ptx3-LECs.

Recently developed algorithms have made important advances in solving the complex problem of integrating different datasets [85; 86; 95]. As shown here they not only perform well in combining replicate samples by removing “batch effects”, but can also identify and map similar subsets of cells across species barriers. Mutual nearest neighbors and CCA algorithms using conserved genes not only mapped human LEC subsets to mouse counterparts, but also uncovered species-specific LN LEC subset compositions, revealing a greater diversity of SCS ceiling LEC in human. We can look forward to continuing advances in computational approaches to integrating and mining scRNA-seq data. Particularly exciting is the power of trajectory inference to recapitulate or predict the organization of endothelial cells within the complex vascular networks. While trajectory analysis has been shown to model sequences of cell phenotypes in development (“pseudotime”) [29], our results both in lymphatics (here) and blood vascular EC studies [30] show that computed alignments of cells in trajectory space reflect the tissue architecture (spatial organization) and physical relationships between endothelial cells *in situ* with surprising faithfulness. This correspondence implies, as an approximation, that endothelial cell phenotypes progress in a gradual and orderly fashion within the linear arrangements (as in vascular tubes) or sheets (as in the SCS and sinus-lining LECs in the LN) that make up the endothelium. In essence, trajectories thus provide a computational roadmap for mapping gene expression to the vascular endothelium. The results underscore the diversity of endothelial cells as a continuum, punctuated by concentrations of particular phenotypes or niches that are identified as “subsets”. Whether the progression of phenotypes and zonation among LN LECs reflects malleable LEC responses to local niche factors, or instead retention of a programmed developmental response, or both, remains to be determined.

Our studies here demonstrate the power of single-cell profiling to illuminate the biology of the vascular endothelium, and the promise it holds to revolutionize our understanding of conserved and species-specific regulation of the vasculature and its responses in physiology and human disease.

## Material and methods

### Mice

Male and female 6- to 8-week-old BALB/cJ or 8- to 12-week-old C57BL/6J peripheral LNs (inguinal, axillary and brachial) (processed by 10x Genomics workflow), or 20-week-old *Prox1-GFP*/C57BL/6J [32] female inguinal LNs (processed by SMART-seq2 workflow) were used for scRNA-seq. Mice were bred and maintained in the animal facilities at Veterans Affairs Palo Alto Health Care System, accredited by the Association for Assessment and Accreditation of Laboratory Animal. In addition, mice were held at a Specific Pathogen Free (SPF) facility Uppsala University, and experimental procedures were approved by the local animal ethics committee at the Uppsala County Court (6009/17).

### Immune challenge

BALB/cJ mice were subjected to cutaneous Oxazolone (Oxa) challenge by applying 5% 4-Ethoxymethylene-2-phenyl-2-oxazolin-5-one (Sigma-Aldrich) in acetone and olive oil topical to the skin as described [74]. Axillary, brachial, and inguinal draining LNs were harvested 48 hours after immunization.

### Tissue dissociation and single cell profiling

Cell isolation for 10x: Single-cell suspensions of total EC were generated as previously described [96]. For each group, axillary, inguinal and brachial LNs from 25-30 male and female BALB/cJ or C57BL/6J mice were combined, minced, washed with Hanks’ Balanced Salt solution, and dissociated for 30 min at 37 °C with gentle rocking in HBSS (with calcium and magnesium) medium containing 0.2 mg/ml Collagenase P, 0.8 mg/ml Dispase II and 0.01 mg/ml DNase I (Sigma-Aldrich) (adapted from [13]). Hematopoietic cells were depleted with anti-CD45 mouse MicroBeads according to the manufacture’s protocol (Miltenyi) and stained with anti-CD31 BV605 (clone 390) and anti-Podoplanin PE-Cy7 (clone 8.1.1) antibodies, as well as dump antibodies consisting of anti-CD45 (clone 30-F11), anti-EpCAM (clone G8.8), anti-TER119 (clone TER-119), anti-CD11a (clone H155-78) and anti-CD11b (clone M1/70). Total EC (lin-CD31+) were sorted into 100% fetal bovine serum using FACS Aria III (BD Biosciences; 100 um nozzle; ∼2500 cells/second), washed, and immediately processed to generate scRNA-seq library using Chromium Single Cell 3’ Library and Gel Bead Kit v2 (10x Genomics) according to manufacturer’s instructions. Libraries were sequenced with NextSeq 500 (Illumina) using 150 cycles high output V2 kit (Read 1: 26 bp, Read 2: 98 bp) at the Stanford Functional Genomics Facility.

Cell isolation for SMART-seq2: Inguinal LN digests from *Prox1-GFP* mice [32], depleted for hematopoietic cells as described above, were stained with anti-CD31 PE-Cy7 (clone 390), anti-Podoplanin AF660 (clone eBio8.11) antibodies (Thermo Fisher Scientific). Dump channel included anti-mouse TER-119 eFluor450 (clone Ter119), anti-mouse CD45 eFluor450 (clone 30-F11), anti-mouse CD11b eFluor450 (clone M1/70) and dead cell staining SYTOX™ Blue (Thermo Fisher Scientific). Triple positive live cells (GFP+ Pdpn+ CD31+) (LECs) were gated and sorted on a BD FACSAria III (BD Biosciences) (100 um nozzle, 20 psi) as single cells into a 384 well plate with lysis buffer. Single cell libraries were prepared as described and sequenced on a HiSeq2500 [97].

## Single-cell RNA-seq data analysis

10x Genomics: Read alignment and quality control were performed using the 10x Genomics Cell Ranger (v3.0.2) and the mm10 reference genome. Loupe Cell Browser (v3.1.1; 10x Genomics) was used to manually gate on LEC (Pdpn+ CD31+) for downstream analysis.

SMART-seq2: After lane demultiplexing, SMART-seq2 based FASTQ files were trimmed with Trim Galore (v0.4.4) followed by alignment of the data to the mouse reference genome (mm10-GRCm38) using TopHat (v2.1.1) and bowtie2 (v2.2.6.0). PCR duplicates were removed using SAMtools (v0.1.18) Counting of fragments aligning per gene was done using the *featurecounts* function of the Subread package (v1.4.6-p5).

Count data were processed with the Seurat package (v3.1.0) [95; 98]. For quality control, genes that were expressed in fewer than 3 cells and cells that expressed fewer than 100 genes were excluded from analysis. Raw counts were log normalized, and 2000 most variable genes were identified based on a variance stabilizing transformation. Variable gene sets were used to align multiple datasets for joint analysis, using the Canonical Correlation Analysis (CCA) method within the ‘FindIntegrationAnchors’ and ‘IntegrateData’ functions of the Seurat package. Principal Component Analysis (PCA) dimensionality reduction was performed using the variable gene sets. Cell clusters were determined using a Shared Nearest Neighbor (SNN) modularity optimization-based clustering algorithm of the Seurat ‘FindClusters’ function, and were visualized with t-distributed Stochastic Neighbor Embedding (tSNE) [99] or Uniform Manifold Approximation and Projection (UMAP) [100]. Contaminating pericyte, immune and blood endothelial cells were removed by supervised gating on the tSNE plot. To recover gene-gene relationships that are lost due to dropouts, we imputed missing gene expression data from log normalized count data using an in-house customization (https://github.com/kbrulois/magicBatch) of the MAGIC (Markov Affinity-based Graph Imputation of Cells) algorithm with optimized parameters (t = 2, k = 9, ka = 3) [101]. Imputed data were used for visualization of single-cell gene expression in violin plots and heatmaps [102], as well as for trajectory analyses.

### Nearest neighbor alignments in trajectory space

tSpace was used to model the nearest neighbor relationships as well as transitional zones between LEC subsets [29]. Batch effects were removed using the ‘fastMNN’ function of the batchelor package (v1.0.1) [103] and variable genes were identified using the scran package [104]. For mouse LEC, we used tSpace with default parameters (T = 100, K = 20, 5 graphs, 20 way points, Euclidean distance metric) and imputed expression values of the top 800 variable genes (Figures 1C and 5A, right panel) or batch-corrected low-dimensional coordinates (top 50 coordinates) (Figure 5A, left panel). The trajectory matrices were visualized in low dimensional space using PCA (Figures 1C and 5A, right panel) or UMAP (Figure 5A, left panel) within the tSpace package. For combined human LEC samples, imputed data of LEC from 6 LNs were batch-corrected as above. Twenty PCs from the 1000 variable genes were calculated, the loadings were adjusted to a minimum value of 0 by addition of a constant and used as input to the tSpace algorithm (T = 200, K = 25, 5 graphs, 10 way points, cosine distance metric).

### Differential gene expression and gene enrichment analysis

Differential gene expression analysis was performed using the negative binomial generalized linear model within the ‘FindMarkers’ function of Seurat on log normalized count data (*p* < 0.01, fold change > 1.2). Only upregulated DEGs were considered, unless otherwise specified. Gene enrichment analysis of DEGs with Gene Ontology[105; 106], Kyoto Encyclopedia of Genes and Genomes [107; 108], and BioPlanet [109] databases was performed using Enrichr [110; 111].

### Correlation analysis

To determine similarities between LEC subsets, mean gene expression values were calculated for each subset. The top 1000 most variable genes across all subsets were used for pairwise Pearson correlation analysis. Hierarchical clustering was performed using the average linkage method.

### Cross species single-cell transcriptome analysis

Human HNLN1 dataset from [26] was used for the integrated LEC profiling. Human gene names were converted to their mouse homologs using the biomaRt package (v2.40.4) [112]. scRNA-seq datasets were integrated in Seurat for unsupervised clustering. The top 100 most upregulated DEGs for each subset, as ranked by the fold change of gene expression in the subset relative to other LEC combined, were determined for human LEC and separately for mouse. An overlap score was defined as the number of DEGs common to one human subset and one mouse subset divided by the number of genes expected to be shared by the two subsets by chance. For each pair of mouse subset and human subsets, overlap score = n / (A × B / N), where n = number of observed overlapping genes between the top 100 DEGs of the human subset and the mouse subset, A = number of DEGs considered in the mouse subset (100), B = number of DEGs considered in the human subset with mouse homologs, and N = total number of genes detected in both mouse and human LEC (13458).

### Reanalysis of human LEC datasets

For the combined analysis of human LEC samples [26], imputed gene expression data were batch-corrected and used for trajectory analysis as above. LEC in each of the additional 5 human samples were classified by correspondence to the index human subsets that mapped with mouse subsets as follows: Reference subset mean gene expression was generated from the index HNLN1 human dataset, using core cells of each major subset that cross-mapped with mouse subsets, as well as manually gated bridge cells that link fLEC and cLEC in tSpace projections. Cells in the other human samples were classified by Pearson correlation using the 1000 most variable genes in the reference set.

### Immunostaining of mouse LNs

Inguinal and popliteal mouse LNs were harvested from *Prox1*-GFP mice [32] and fixed in 0.8% paraformaldehyde (PFA) for 12h at 4°C. After fixation, the LNs were placed in sucrose: 25% for 2 days, 50% for 2 days before embedding in OCT media (HistoLab), then snap frozen on dry ice. Frozen tissues were cryo-sectioned at a thickness of 8µm and stored at −80°C. For fresh frozen tissue, LNs were harvested from wild-type C57BL/6 mice and cleaned in ice cold PBS, embedded in OCT media and snap frozen on dry ice. For immunostaining, the sections were hydrated in Phosphate-buffered saline (PBS) and blocked with 10% donkey serum (Sigma) diluted in PBS for 20 min. After blocking, the sections were incubated with primary antibodies diluted in blocking buffer overnight at 4°C. Thereafter the sections were washed in PBS with 0.1% TritonX100 (Sigma) (PBSTX) and incubated with secondary antibodies diluted in PBSTX for 1h at RT. The sections were counterstained with 4′,6-diamidino-2-phenylindole (DAPI) following additional washing in PBSTX and mounted in ProLong Gold Antifade Mountant (Thermofisher Scientific).

### Immunostaining of human LNs

Human head and neck tumor-free LNs from cancer patients were received from the hospital immediately after the surgery, and embedded in OCT compound (Sigma) and frozen on dry ice. The collection was done under the license ETMK: 132/2016. A written informed consent was obtained from each individual donating tissue. The samples were kept anonymous and used with the permission of the Ethical Committee of Turku University Hospital. The LNs were sectioned at a thickness of 6 µm with a cryostat, and fixed with acetone at −20 °C. The sections were incubated with 10% FCS for blocking, incubated with the primary antibodies diluted in 0.5% BSA in PBS overnight at 4 °C. Thereafter, they were incubated with the secondary antibodies for 2 hr at room temperature. Sections were washed with PBS and mounted with ProLong Gold Antifade Mounting medium with DAPI (Thermofisher Scientific).

### Antibodies

Primary antibodies for mouse antigens: anti-eGFP (Abcam, clone 7.1 and 13.1), anti-MAdCAM-1 (eBioscience, MECA-367), anti-Lyve-1 (ReliaTech, 103-PA50), anti-PD-L1 (BioLegend, 10F.9G2), anti-Marco (Bio-Rad, ED31) anti-B220/CD45R (ebiosciences, RA3-6B2). Primary antibodies for human antigens: anti-PROX-1 (R and D, AF2727), anti-LYVE-1 (Reliatech, 102-PA50), anti-CLEC4M (R and D, MAB162), anti-MARCO (Sigma, HPA063793), anti-CD36 (Abcam, ab17044). Secondary antibodies: donkey-anti-chicken AF488 (Jackson ImmunoResearch), donkey anti-rabbit Cy3 (Jackson ImmunoResearch), donkey anti-rabbit AF647 (Invitrogen), goat anti-rabbit AF546 (Invitrogen), donkey anti-mouse Cy3 (Jackson ImmunoResearch), donkey anti-mouse AF647 (invitrogen), donkey anti-goat AF488, AF555 and AF594 (Invitrogen), donkey anti-rat AF488 and AF647 (Jackson ImmunoResearch).

### RNA in situ hybridization

Inguinal mouse LNs were harvested from wild-type C57BL/6 mice following fixation with 2% PFA by heart perfusion. The tissues were embedded in OCT and snap frozen on dry ice. The tissue were cryosectioned at a thickness of 14µm, dried at −20°C for 1 hr and stored at −80°C. In situ hybridization (ISH) was performed using RNAscope Multiplex Fluorescent kit according to the manufacturer’s instructions (Advanced Cell Diagnostics). Briefly, the sections were fixed in ice-cold 4% PFA for 15 min, rinsed in PBS and dehydrated with increasing concentrations of ethanol: 50%, 70% and absolute ethanol for 5 min each. The sections were dried at RT and treated with protease IV for 15 min and rinsed in PBS. Thereafter the sections were incubated with the mouse probes: *Claudin-5-C3, Stabilin-1-C1, Bmp2-C2, Bmp4-C2* in ACD HybEZ II hybridization system (Advanced Cell Diagnostics) at 40°C for 2h. The remainder of the assay protocol was implemented following manufacturer’s statement. The sections were then counterstained with DAPI.

### Imaging

Images were obtained with Vectra Polaris™ Automated Quantitative Pathology Imaging System (Akoya Biosciences) and LSM 700 or LSM 880 confocal microscopes (Zeiss). Confocal objectives: Air objective plan apo N/A 0.45 (10x magnification); air objective plan apo N/A 0.8 (20x magnification); air objective plan apo N/A 0.95 (40x magnification). Analyses were performed with ImageJ software.

### Data availability

Data will be deposited in the NCBI Gene Expression Omnibus database. A searchable database with gene expression visualization for human and mouse datasets will also be available in the format of a website.

### Author contributions

MX performed bioinformatic analyses with contribution from KB. RG, AT, KB, TB, and SN performed experiments. RG and TB performed the mouse image analysis and illustration. AT performed human image analysis. MX, RG, MHU and ECB wrote and edited the manuscript. JP contributed to the content. KB, TB, DD, AT and SJ provided critical input and edited the manuscript. MV contributed advice. MHU and ECB directed the study.

### Funding

This work was supported by the Swedish Research Council (2016-02492), Swedish Cancer Foundation (2017/759) Kjell and Märta Beijer Foundation and Malin and Lennart Philipsson Foundation to MHU; and by NIH grants R01-AI130471 and R37-AI047822, MERIT award I01 BX-002919 from the Dept of Veterans Affairs, and pilot awards under ITI Seed 122C158 and CCSB grant U54-CA209971 to ECB. KB was supported by NIH F32 CA200103, and SN by the Swedish Society for Medical Research and Stanford Dean’s Fellowship. RG was supported by GA Johanssons foundation. AT was supported by the Academy of Finland.

## Acknowledgments

We would like to thank Sonja Gustafsson, Jianping Liu and Byambajav Buyandelger (Single cell core unit, ICMC, Karolinska Institute) for preparation and sequencing of the SMART-seq2 libraries and Liqun He for initial analysis. We thank Theresa Dinh, Anusha Rajaraman, Romain Ballet, Yu Zhu and Wing Lam for sample processing, Nicole Lazarus for technical advice, Dhananjay Wagh for 10x single cell sequencing, and Riika Pietlä for technical support and advice on RNAscope.

## Supplementary Figure Legends

**Figure S1:**
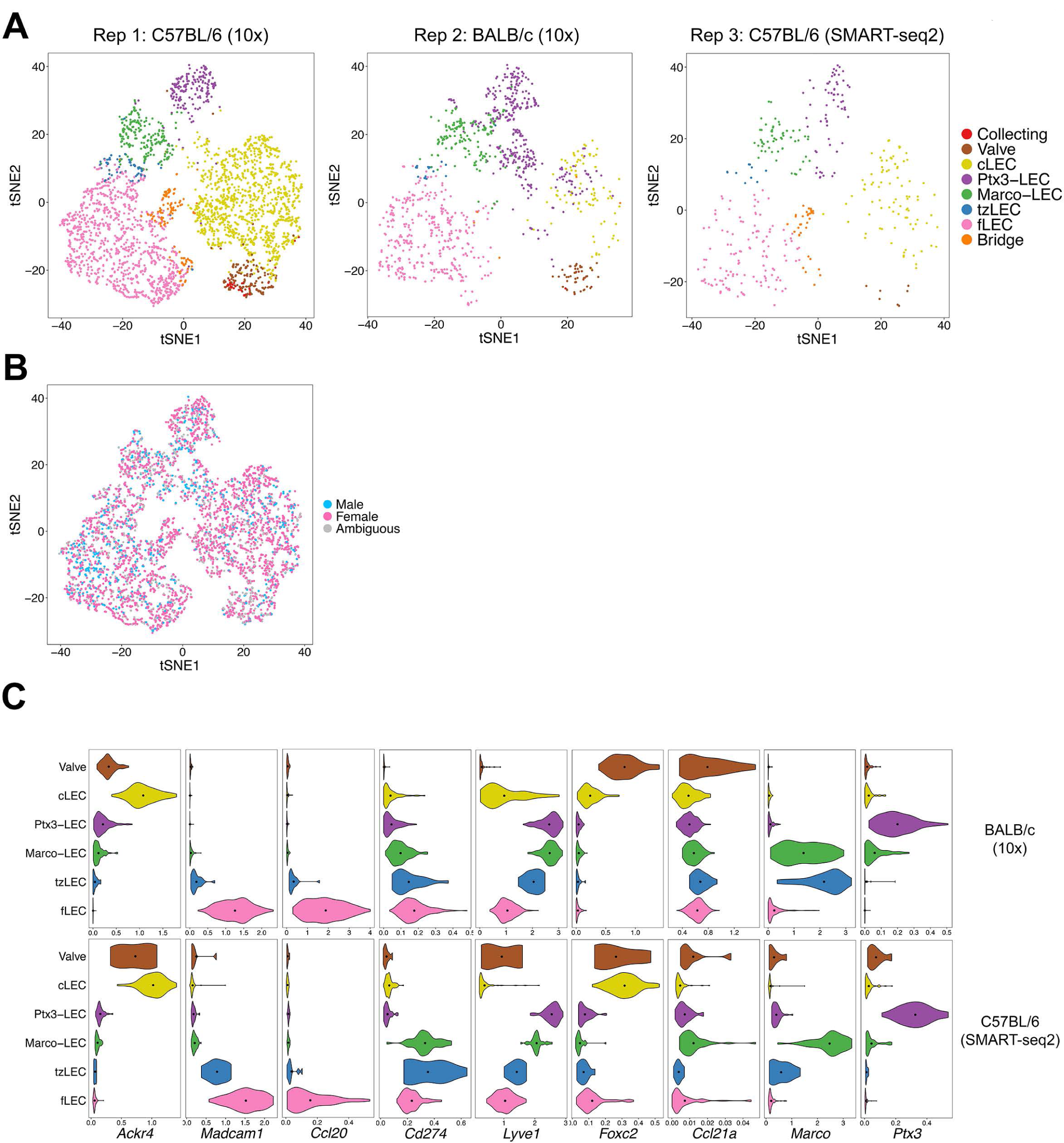
Mouse LN LEC subsets in biological replicates. **(A)** tSNE plots of LEC from three individual biological replicates, colored by subset. **(B)** tSNE plot of LEC from all replicates, colored by sex. **(C)** Expression of subset defining genes in BALB/c (10x) and C57BL/6 (SMART-seq2) mice. Dots indicate mean log-normalized transcript count.

**Figure S2:**
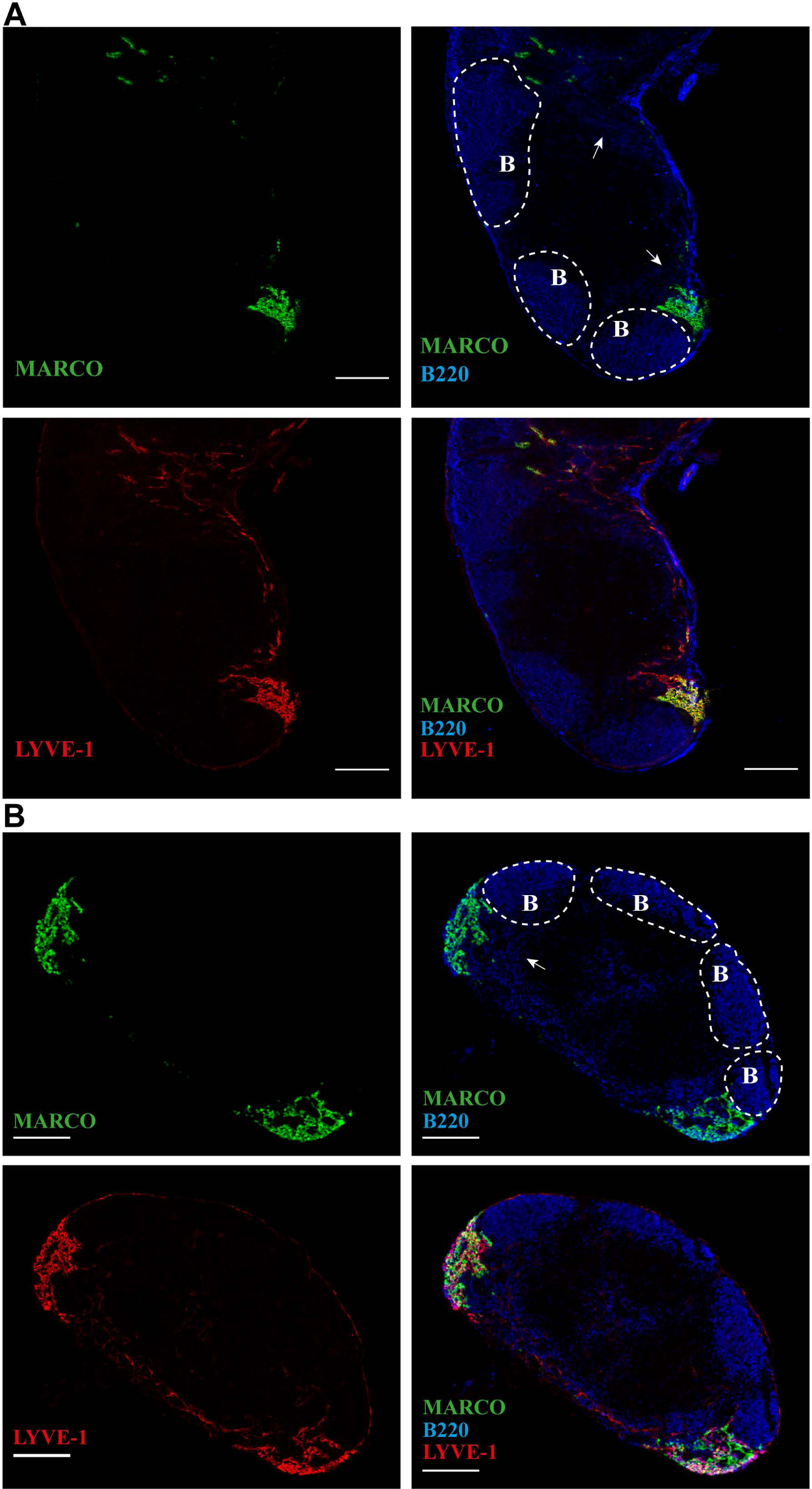
MARCO+ LECs are close to B-cell follicular area. B220 (blue), LYVE-1 (red) and MARCO (green) in inguinal **(A)** and popliteal **(B)** LN sections of wild type mice. B-cell follicular areas are indicated (white dashed lines) and scattered B-lineage plasma cells in the medulla (white arrows). Data are representative of three or more independent experiments. Scale bar = 200µm.

**Figure S3:**
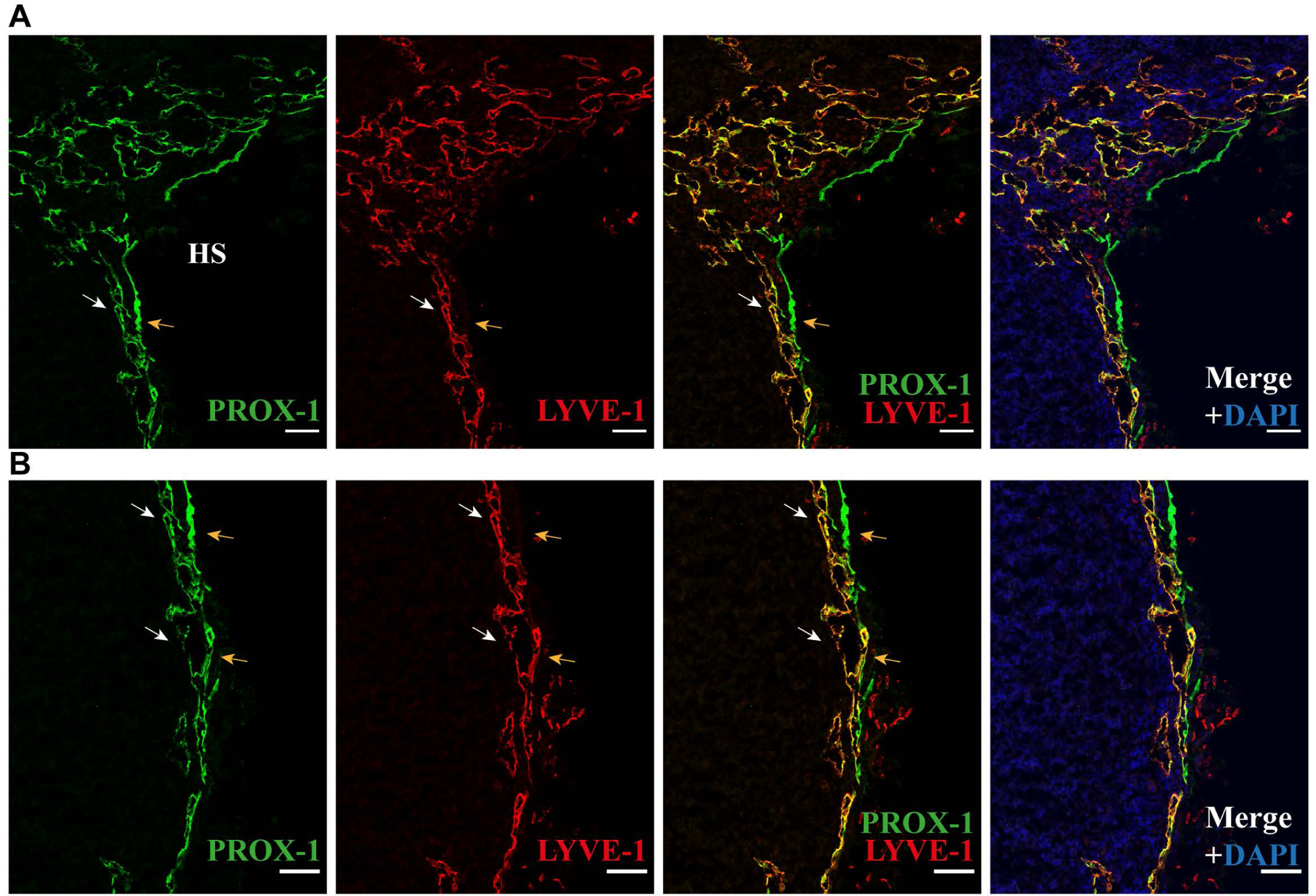
Interaction between Ptx3-LECs and cLECs in peri-hilar sinuses. Inguinal LNs from *Prox-1-GFP* transgenic mouse stained for GFP (PROX-1) (green) and LYVE-1 (red), counterstained with DAPI (blue). HS = hilus. LYVE-1+ LECs (Ptx3-LEC area) are indicated with white arrows and LYVE-1-cLECs are indicated with orange arrows. Data are representative of three or more independent experiments. Scale bar = 50µm.

**Figure S4:**
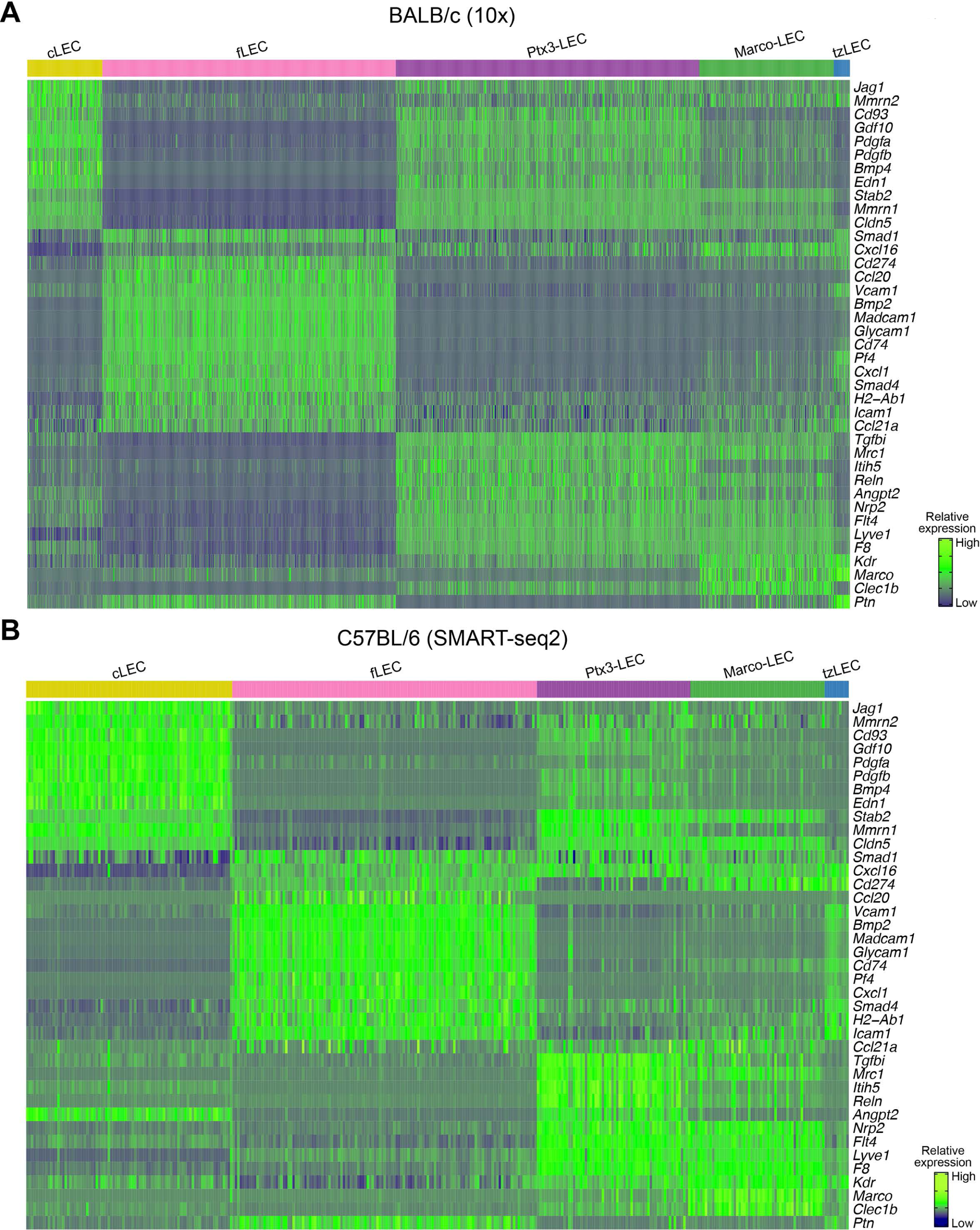
Heatmaps of LEC subset DEGs in biological replicates. Heatmaps of select DEGs in LECs of BALB/c (10x) and C57BL/6 (SMART-seq2) mice. Values are imputed log counts (row scaled).

**Figure S5:**
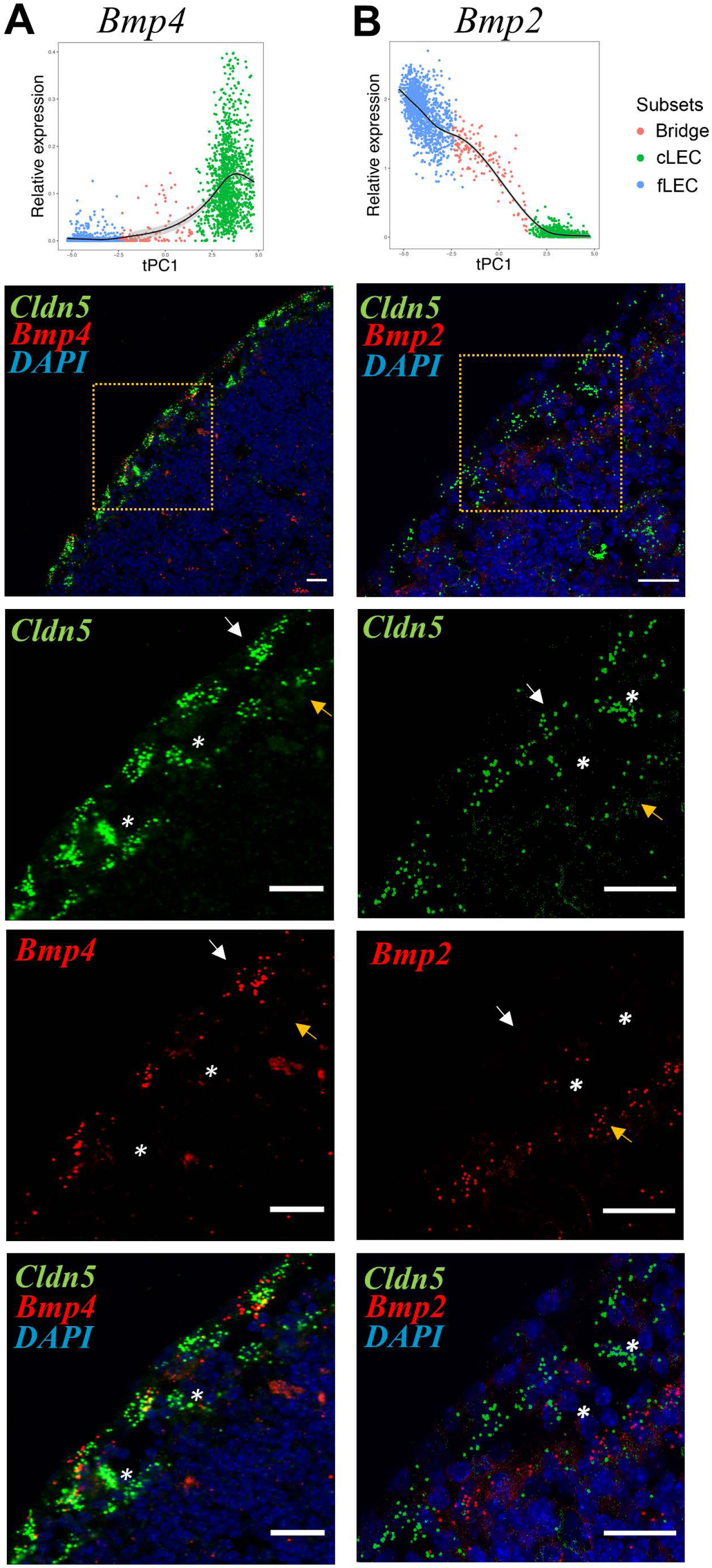
Bmp4 and *Bmp2* differentiate cLECs from fLECs respectively and SCS bridging cells. *In situ* hybridization (RNAscope-ISH) of mouse inguinal LNs. **(A)** mRNA detection of *Claudin-5* (*Cldn5*) (green) and *Bmp4* (red) with fluorescent probes. ROI inset (orange dotted box) shown below. **(B)** mRNA detection of *Claudin-5* (*Cldn5*) (green) and *Bmp2* (red) with fluorescent probes. ROI inset (orage dotted box) shown below. Counterstain is DAPI (blue). The ceiling lymphatic endothelial cells (cLECs) (white arrows) and the lymphatics endothelium lining the floor (fLECs) (orange arrows) are indicated. Bridges are indicated with white stars. scRNA-seq expression across cLEC, bridge and fLEC populations is shown above. Outliers are not shown. Scale bar = 20µm.

**Figure S6:**
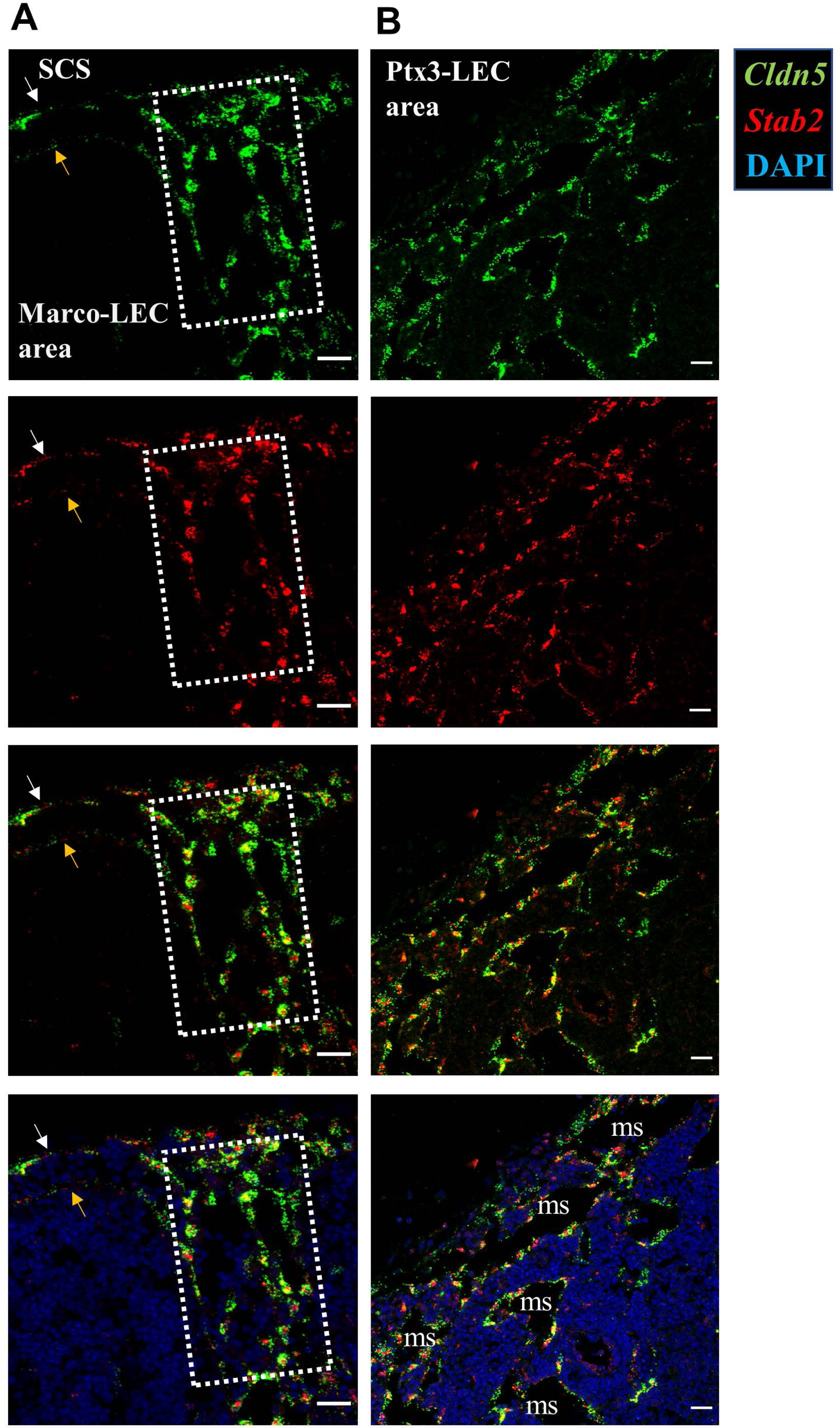
Low *Stabilin-2* expression differentiates fLECs from medullary populations and cLECs. *In situ* hybridization (RNAscope-ISH) of mouse inguinal LNs. Detection of *Cldn5* (green) and *Stab2* (red) mRNA with fluorescent probes, counterstained with DAPI (blue). **(A)** Subcapsular sinus area; cLECs (white arrows) and fLECs (orange arrows) populations are indicated. Peri-follicular medulla (corresponding to Marco-LECs) is outlined with white dotted rectangle. **(B)** Central medulla (peri-hilar) on the efferent (eff) side of the LN (Ptx3-LECs). The medullary sinuses (ms) are indicated. Scale bar = 20µm.

**Figure S7:**
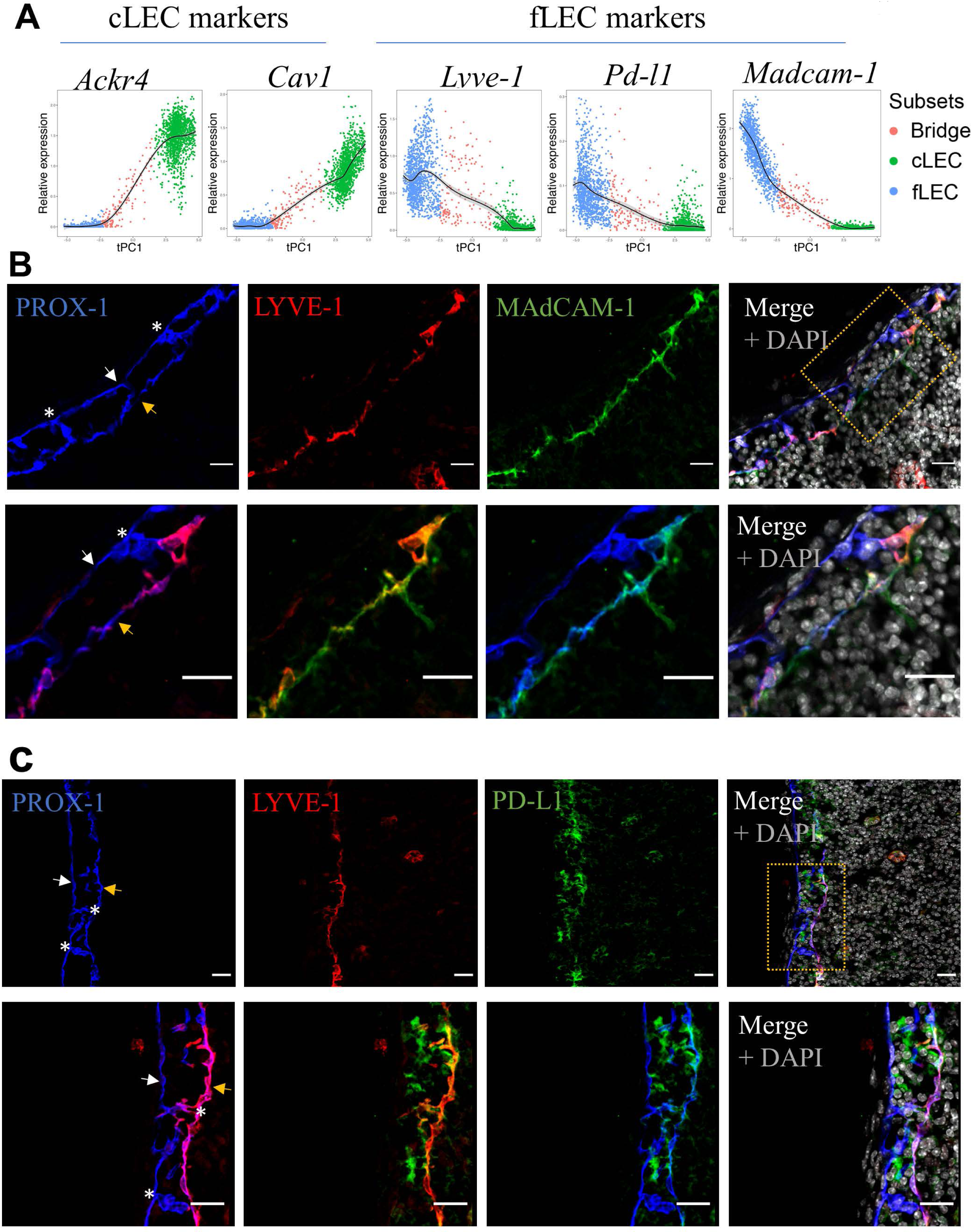
SCS bridging cells. **(A)** scRNA-seq expression of cLEC and fLEC marker genes across cLEC, bridge and fLEC populations. Outliers are not shown. **(B, C)** Immunoreactivity of GFP (Prox-1-GFP) (blue), LYVE-1(red) and MAdCAM-1 **(B)** or PD-L1 **(C)** (green), in inguinal LNs from *Prox-1-GFP* transgenic mice, counterstained with DAPI (gray). Area of insets is shown by orange dotted rectangle. The ceiling lymphatic endothelial cells (cLECs) (white arrows), the lymphatic endothelium lining the floor (fLECs) (orange arrows) and bridge population (white stars) are indicated. Data are representative of three or more independent experiment. Scale bar = 20µm.

**Figure S8:**
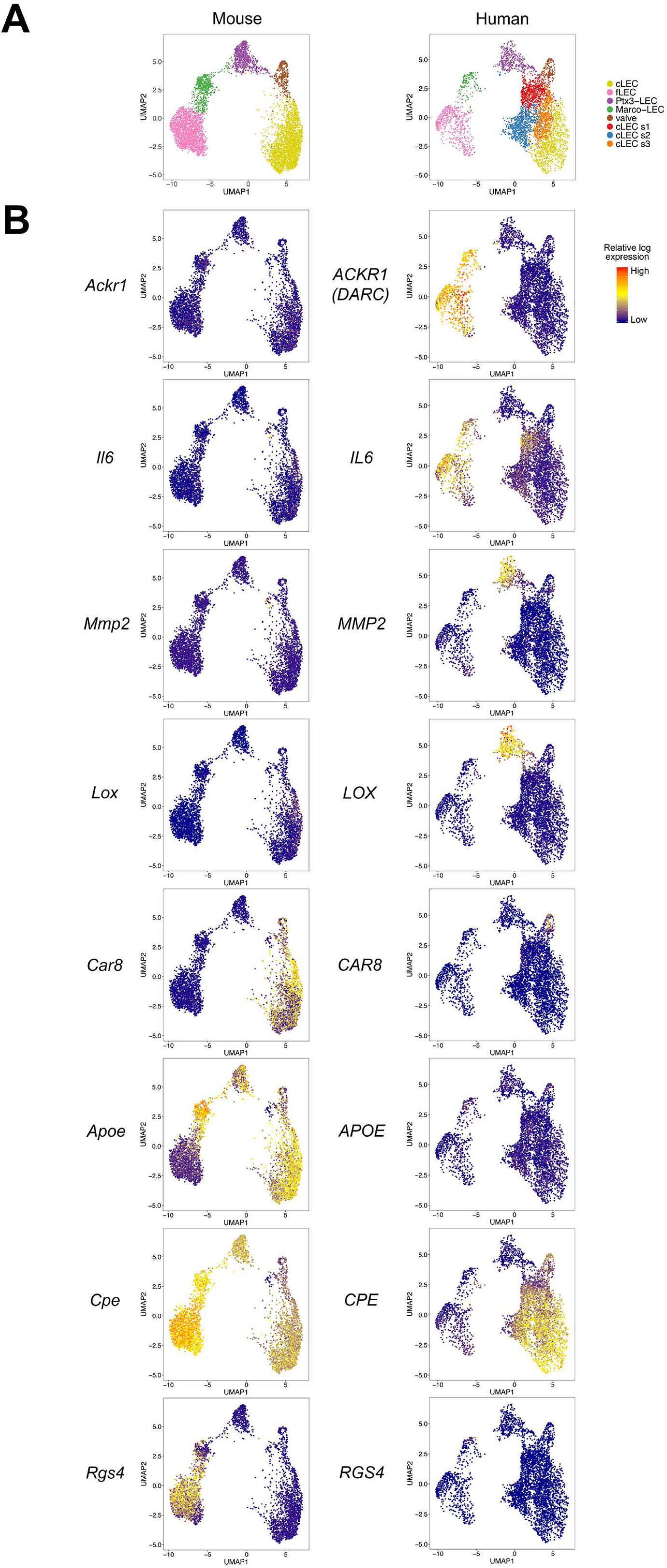
Illustration of differential gene expression patterns in mouse and human. **(A)** UMAP of aligned mouse and human LEC, colored by subset (reproduced from Figure 6A). **(B)** Expression pattern of indicated genes, projected on UMAP plot of mouse (left) and human (right) LN LECs. Values are imputed log counts.

